# N-glycan chitobiose core biosynthesis by Agl24 strengthens the hypothesis of an archaeal origin of the eukaryal N-glycosylation

**DOI:** 10.1101/2021.01.19.427365

**Authors:** Benjamin H. Meyer, Ben A. Wagstaff, Panagiotis S. Adam, Sonja-Verena Albers, Helge C. Dorfmueller

**Author notes:** Correspondence to B. H. Meyer, and H. C. Dorfmueller. **Cover Art:** 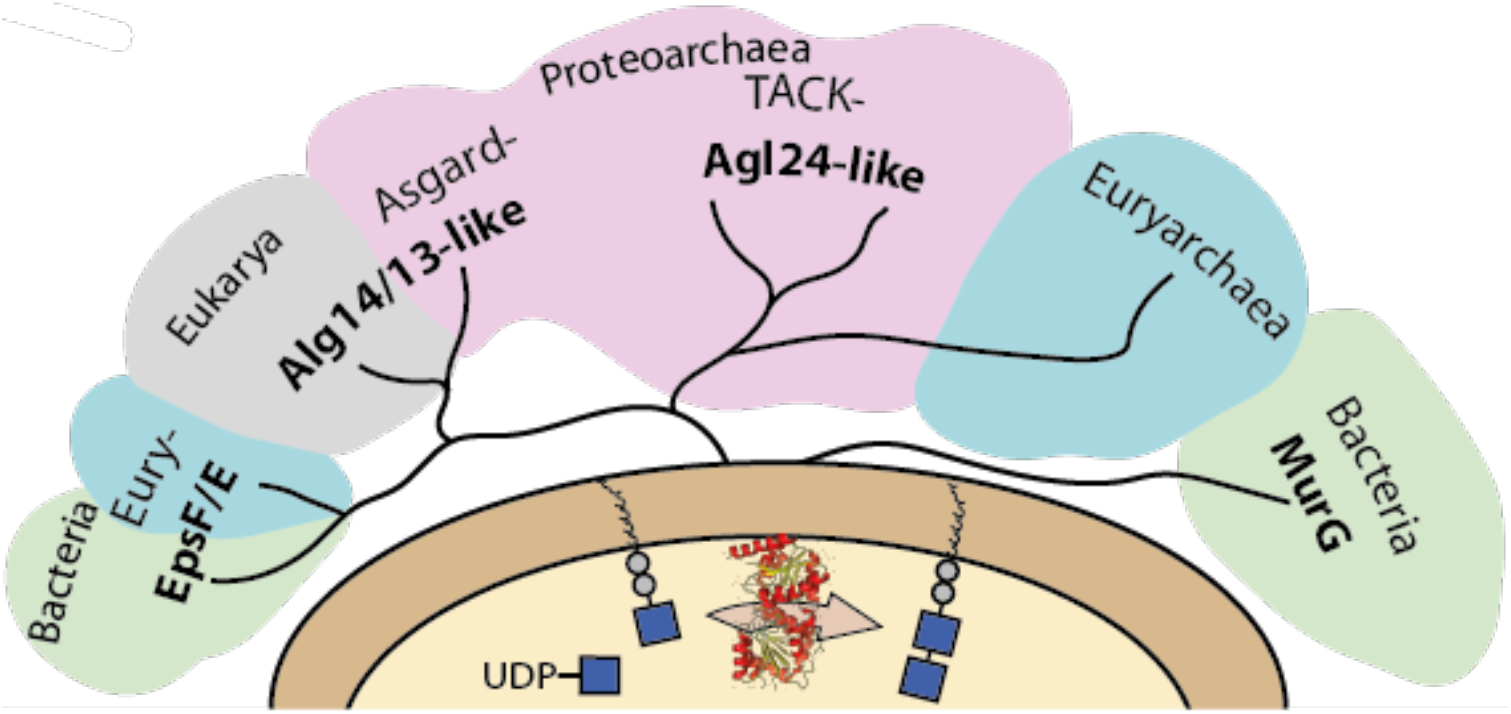.

## Abstract

Protein N-glycosylation is the most common posttranslational modifications found in all three domains of life. The crenarchaeal N-glycosylation begins with the synthesis of a lipid-linked chitobiose core structure, identical to that in eukaryotes. Here, we report the identification of a thermostable archaeal beta-1,4-N-acetylglucosaminyltransferase, named archaeal glycosylation enzyme 24 (Agl24), responsible for the synthesis of the N-glycan chitobiose core. Biochemical characterization confirmed the function as an inverting β-D-GlcNAc-(1→4)-α-D-GlcNAc-diphosphodolichol glycosyltransferase. Substitution of a conserved histidine residue, found also in the eukaryotic and bacterial homologs, demonstrated its functional importance for Agl24. Furthermore, bioinformatics and structural modeling revealed strong similarities between Agl24 and both the eukaryotic Alg14/13 and a distant relation to the bacterial MurG, which catalyze the identical or a similar process, respectively. Our data, complemented by phylogenetic analysis of Alg13 and Alg14, revealed similar sequences in Asgardarchaeota, further supporting the hypothesis that the Alg13/14 homologs in eukaryotes have been acquired during eukaryogenesis.

**Highlights:**

- First identification and characterization of a thermostable β-D-GlcNAc-(1→4)-α-D-GlcNAc-diphosphodolichol glycosyltransferase (GT family 28) in Archaea.
- A highly conserved histidine, within a GGH motif in Agl24, Alg14, and MurG, is essential for function of Agl24.
- Agl24-like homologs are broadly distributed among Archaea.
- The eukaryotic Alg13 and Alg14 are closely related to the Asgard homologs, suggesting their acquisition during eukaryogenesis.

## Introduction

Asparagine (N)-linked glycosylation is one of the commonest co- and posttranslational protein modifications found in all three domains of life (Larkin and Imperiali, 2011, Jarrell et al., 2014, Nothaft and Szymanski, 2010) In Eukarya, N-glycosylation is an essential process and is evolutionarily highly conserved from yeast to humans (Lehle et al., 2006). It is estimated that more than half of all eukaryotic proteins are glycoproteins (Apweiler et al., 1999, Zielinska et al., 2010). N-glycosylation is required for correct protein folding and stability, intra- and extracellular recognition of protein targets and for enzyme activity (Caramelo and Parodi, 2007, Helenius and Aebi, 2004, Varki, 1993). Therefore, the biological functions of protein glycosylation range from relatively minor to crucial for survival of an organism. As such, congenital disorders of glycosylation in humans are often lethal or cause severe phenotypes and diseases ranging from intellectual disability, growth retardation, cardiac anomalies, and early death (Reily et al., 2019). Although it was long thought that N-glycosylation was restricted to Eukarya, N-glycosylation pathways have now been characterized in all three domains of life. Bacterial N-glycosylation seems to be restricted to delta/epsilon proteobacteria (Nothaft and Szymanski, 2013, Whitfield et al., 2017). In contrast, all sequenced Archaea possess the N-glycan oligosaccharyltransferase AglB, with the exceptions of *Aeropyrum pernix* and *Methanopyrus* species (Jarrell et al., 2014, Nikolayev et al., 2020). The common feature of N-glycosylation is the assembly of an oligosaccharide onto a lipid by the action of specific glycosyltransferases (GTs), which use either nucleotide- or lipid-activated sugar donors. The assembled lipid-linked oligosaccharide (LLO) is flipped across the cytoplasmic/ER membrane. The glycan is then either enlarged or directly transferred onto a target protein by an oligosaccharyltransferase (OST). Target proteins contain a specific asparagine residue found within the conserved sequon (N-X-S/T in Eukarya and Archaea or D/E-X_1_-N-X_2_-S/T in Bacteria, where X is not proline) recognized by the OST.

Archaeal N-glycosylation displays a mosaic of bacterial and eukaryal features. Like Bacteria, Archaea use only a single OST subunit, whereas Eukarya require an OST complex of nine subunits, to transfer the LLO onto the target protein. In contrast, Archaea use the shorter version of the recognition sequon (N-X-S/T) as well as dolichol-phosphate (Dol) as lipid, as in Eukarya, whereas Bacteria use undecaprenyl-phosphate. Archaea also display unique N-glycosylation features, *e.g.* the assembly of the N-glycan onto Dol-phosphate in the Euryarchaeota, while the Crenarchaeota use Dol-pyrophosphate as in eukaryotes (Taguchi et al., 2016).

Currently, more than 15 different archaeal N-glycan structures have been characterized (Jarrell et al., 2014, Albers et al., 2017). These archaeal N-glycans display unique structures and sugar compositions and based on the limited characterization of archaeal N-glycans, it is plausible to assume that Archaea possess an enormous range of structural and composition diverse N-glycans. However, closely related species *e.g.,* of the thermophilic crenarchaeal Sulfolobales family, retain a conserved N-glycan core structure but differentiated by their terminal sugar residues. The glycosylation pathway of *Sulfolobus acidocaldarius*, belonging to Sulfolobales, has been partially characterized. Secreted proteins, such as the cell wall surface-layer (S-layer) protein and motility filament forming protein ArlB, are N-glycosylated with a tribranched hexasaccharide. This N-glycan has the composition: (Glc-β1-4-sulfoquinovoseβ-1-3)(Man α-1-6)(Man α-1-4)GlcNAc-β1-4-GlcNAc-β-Asp (Peyfoon et al., 2010, Zahringer et al., 2000). Interestingly, the N-glycan core structure is composed of a chitobiose (GlcNAc-β1-4-GlcNAc), which is identical to the core structure of all eukaryal N-glycans. Members of the Sulfolobales family are the only Archaea reported to display this eukaryal N-linked chitobiose core structure (van Wolferen et al., 2020, Palmieri et al., 2013). Based on this identical N-glycan core structure, it has been suggested that the *Sulfolobus* N-glycan biosynthesis pathway may be similar to the eukaryal process (Meyer and Albers, 2013). Indeed, a homolog of the yeast Alg7 enzyme, which initiates eukaryal N-glycosylation by transferring a GlcNAc-P from UDP-GlcNAc to Dol-P to generate the precursor Dol-PP-GlcNAc, was identified in *S. acidocaldarius* as AlgH (Meyer et al., 2017). This AglH showed not only a similar topology and highly conserved amino acid motifs to Alg7, but also functionally complemented a conditional yeast Alg7 knockout (Meyer et al., 2017). This is a rare example of cross-domain complementation that further strengthens the hypothesis of a common origin of eukaryotic and archaeal N-glycosylation.

Here we report the identification of the β-1,4-N-acetylglucosamine-transferase Agl24, which shows conserved amino acid motifs and apparent structural similarities with the eukaryal Alg13/14 and bacterial MurG functional homologs. Activity assays, along with HPLC, MALDI, and NMR analyses confirmed the function of Agl24 as catalyzing the second N-glycan biosynthesis step by generating the lipid-linked chitobiose core. Our extensive bioinformatics analyses support the hypothesis that the eukaryotic N-glycosylation pathway originates from an archaeal ancestor.

## Results

### Identification of a Dol-PP-GlcNAc UDP-GlcNAc GlcNAc GT candidate in *S. acidocaldarius*

To identify the glycosyltransferase that conducts this first committed synthetic step in the N-glycosylation process in *S. acidocaldarius*, we searched for similar biosynthesis processes to identify potential homologs. The bacterial enzyme MurG transfers a GlcNAc residue from UDP-GlcNAc to the C4 hydroxyl group of MurNAc-(pentapeptide)-PP-undecaprenol to produce the lipid-linked β-(1,4) disaccharide known as lipid II (Ha et al., 1999, Men et al., 1998). Although MurG does not synthesize chitobiose like *S. acidocaldarius*, the enzyme uses a similar lipid-linked acceptor and an identical sugar donor (UDP-GlcNAc) and therefore presents a valid candidate for evaluation. MurG has been classified by the Carbohydrate Active enZyme database (http://www.cazy.org/) as a family 28 GT(Lombard et al., 2014). Up to now, only members of the archaeal families *Methanobacteriaceae* and *Methanopyraceae*, which synthesize pseudomurein as their cell wall structure (Albers and Meyer, 2011, Meyer and Albers, 2020), were found to have a GT28 family homolog in the CAZy database (Lombard et al., 2014).

Another candidate homolog is the eukaryal protein complex formed by Alg13 and Alg14 that transfers GlcNAc from UDP-GlcNAc to the C4 hydroxyl of GlcNAc-PP-dolichol to produce the β-(1-4) chitobiose core of the eukaryal N-glycan. Alg13 and Alg14 are classified as GT 1 family members but are distantly related to GT28 family enzymes (Lombard et al., 2014). Alg13 and Alg14 form the C- and N-terminal domains, respectively, of a functional heterodimeric GT enzyme (Fig 2); Alg14 contains the acceptor-binding site and the membrane association domain, while Alg13 possesses the UDP-GlcNAc-binding pocket. The interaction of these two enzymes is required for the catalytic activity, as the active site lies in the cavity formed by the two proteins (Gao et al., 2005, Bickel et al., 2005, Chantret et al., 2005).

Although members of the GT 1 family are most prominent in Archaea, until now no GT1 family member has been classified in Sulfolobales. There is no meaningful shared sequence identity between Alg14-13 and MurG and they are subsequently annotated in different GT families. However, both enzymes use UDP-GlcNAc to GlcNAcylate a lipid-linked sugar acceptor and therefore represent valid candidates for a bioinformatics search in *S. acidocaldarius.*

Using the *E. coli* MurG protein sequence (WP_063074721.1; EC 2.4.1.227) as template for a Delta Blast search (Boratyn et al., 2012), various homologs were identified in *S. acidocaldarius* with very low sequence identity of only 10–17% (Table S1). The anchoring subunit Alg14 from *Saccharomyces cerevisiae* S288C (NP_009626.1) was used as bait for a homology search but no protein candidate was identified below the threshold of 0.005. However, when either the catalytic subunit Alg13 from *Saccharomyces cerevisiae* S288C (NP_011468.1; EC 2.4.1.141) or the fusion of both Alg14 and Alg13 sequences was used as a search model, a single Sulfolobales homolog was identified with 17% sequence identity (Table S1). Thus, the most likely candidate for the β-1,4-N-acetylglucosamine transferase was found to be Saci1262 (Uniprot: Q4J9C3), now named Agl24, a hypothetical 327 amino acid protein. In contrast to Euryarchaeota, where the genes of the N-glycosylation enzymes are clustered with *aglB,* clustering of these genes is uncommon in Crenarchaeota (Kaminski et al., 2013, Nikolayev et al., 2020), which makes the identification of GTs involved in the N-glycosylation process more challenging, as in the Sulfolobales. Interestingly, the gene *saci1262 / agl24* is located only eight genes downstream of the OST *aglB in S. acidocaldarius.* (Fig S1).

### Detailed bioinformatics comparison of Agl24 with Alg13/14 and MurG

The overall topology of Agl24 is identical to MurG, which consist of two domain folds separated by a deep cleft corresponding to a GT-B fold structure (Ha et al., 2000, Hu et al., 2003) (Fig 1C). Sequence and domain comparisons of the eukaryal Alg13 and Alg14 to MurG and Agl24 reveal that Alg14 corresponds to the N-terminal part of MurG/Agl24, while Agl13 resembles the C-terminal part of MurG/Agl24 (Fig 1A). Both N-terminal (aa1–152) and C-terminal (aa153–327) parts of Agl24 show very low sequence identity to Alg14 and Alg13, with 15% and 16%, respectively. Interestingly, in lower eukaryotes, such as Leishmania and Trypanosoma, Alg14 and Alg13 enzymes are present as a single protein fusion, similar to *S. acidocaldarius* (Averbeck et al., 2007). The question arises whether the ancestral enzyme was fused and split into two domains during evolution. Strikingly, the opposite domain orientation is found *e.g.,* in *Entamoeba histolytica* (Bickel et al., 2005) and *Dictyostelium discoideum*. An interesting difference in these proteins from the three domains of life is the presence of an N-terminal transmembrane domain (TMD) in most of eukaryal Alg14 proteins (aa 5–25), which is missing in bacterial MurGs and archaeal Agl24 enzymes. The TMD is predicted to facilitate the association of Agl14 with the ER membrane and to participate in recruitment of Dol-PP-GlcNAc to the membrane (Lu et al., 2012). The introduction of this TMD appears to be derived later in the evolution of eukaryotes, and the interaction with the ER or cytoplasmic membrane is not solely dependent on a TMD, as the presence of hydrophobic patches and protein–protein interactions with membrane proteins are also proposed to facilitate this interaction. Indeed, in lower Eukarya, *e.g.* in *D. discoideum* and *E. histolytica,* having the exchanged domain orientation, lacking an N-terminal TMD (Averbeck et al., 2007).

**Fig. 1.**
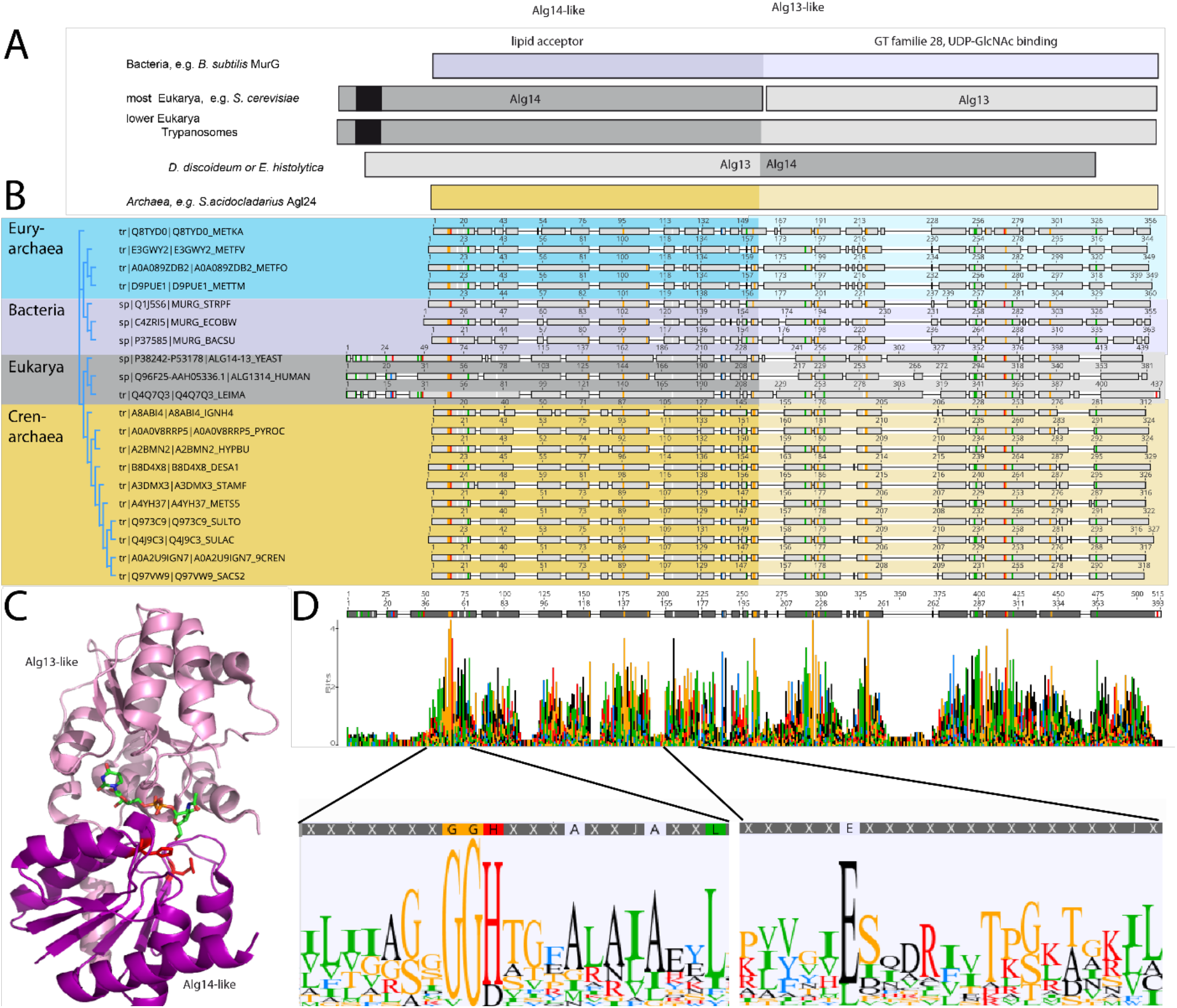
Comparison of the archaeal Agl24, bacterial MurG, and eukaryal Alg14-13. **A)** Simplified comparison of the Agl14-like (darker) or Alg13-like (light color) domain in eukaryal Alg14-13, archaeal Agl24, and bacterial MurG ortologs. Transmembrane domain is depicted in black. **B)** Protein sequences were aligned with the Clustal Omega (Sievers et al., 2018) (for the full alignment see Fig S2). Sequences derived from i) three bacterial MurG: *E.coli* (C4ZRI5), *Streptococcus pyogenes* (Q1J5S6), and *Bacillus subtilis* (P37585); ii) eukaryal Alg14-13: *Leishmania major* (Q4Q7Q3), an artificial Alg14-13 fusion from *Saccharomyces cerevisiae* (P38242-P53178) and *Homo sapiens* (Q96F25-Q9NP73-2); iii) the crenarchaeal Agl24 homologs: *Ignicoccus hospitalis (*A8ABI4), *Pyrodictium occultum* (A0A0V8RRP5), *Hyperthermus butylicus (*A2BMN2), *Desulfurococcus amylolyticus* (B8D4X8), *Staphylothermus marinus (*A3DMX3), *Metallosphaera sedula* (A4YH37), *Sulfurisphaera tokodaii* (Q973C9), *Sulfolobus acidocaldarius* (Q4J9C3), *Acidianus brierleyi* (A0A2U9IGN7), and *Saccharolobus solfataricus (*Q97VW9). Selected sequences from pseudomurein producing Euryarchaea with higher sequence similarity to the MurG are aligned: *Methanopyrus kandleri* (Q8TYD0), *Methanothermus fervidus (*E3GWY2), *Methanobacterium formicicum* (A0A089ZDB2), *Methanothermobacter marburgensis* (D9PUE1). Conserved amino acids (65% threshold) are highlighted in color corresponding to their chemical properties of the amino acids: green polar amino acids (G, S, T, Y, C, Q, N), blue basic (K, R, H), red acidic (D, E), and black hydrophobic amino acids (A, V, L, I, P, W, F, M). End of the Agl14-like domain and start of Alg13-like domains are indicated by the change from dark to light background color. **C)** Ribbon presentation of MurG (PDB: 3s2u) bound to UDP-GlcNAc (sticks, green carbon, blue nitrogen, red oxygen) and orientation towards the cell membrane (bottom). Agl13-like domain (turquoise) and Agl14-like domain (violet) are labelled. Conserved His and Glu residues in Agl14 are shown as red sticks. **D)** Strict consensus (65%) and weblogo from the sequence alignment shown in B). Height of each bar correspond to the observed frequency, highly conserved motifs or amino acids are enlarged below, including conserved motifs G(x)GGH15 (Loop I) and E_114_ (full Weblogo see Fig S2).

The amino acid sequence alignment of the representative bacterial MurG, eukaryal Alg14-13, and archaeal homologs revealed conservation of specific patches and individual residues (Fig 1B and 1D). Despite the low sequence identity, a similar structural topology of Agl24 to Agl14/13 and MurG is not surprising, as GTs in general show only a limited number of structural folds. GTs mainly possess a GT-A or GT-B fold, whereas oligosaccharyltransferases and a number of recently characterized GTs display a GT-C, GT-D, or GT-E folds, respectively (Mestrom et al., 2019). Crystal structures of the bacterial MurG (PDB:3s2U, 1F0K, 1NLM) and the C-terminal half of the eukariotic Alg13 (PDB:2jzc) show typical GT-B structural characteristics (Brown et al., 2013, Ha et al., 2000, Hu et al., 2003, Wang et al., 2008, Raman et al., 2010) (Fig 2). The N-terminal part of Alg13 contains a Rossmann-like fold with a mixed parallel and antiparallel β-sheet rather than the conventional Rossmann fold found in all GT-B enzymes, indicating a unique topology among glycosyltransferases(Wang et al., 2008). The N-terminal helix of Agl14 and MurG is proposed to be the membrane association domain (Chantret et al., 2005, Lu et al., 2011). We have generated an Agl24 structural model to investigate potential features via SWISS-MODEL (Waterhouse et al., 2018) (Fig 2). Previous studies have compared the structures of MurG and Agl13, including a model for Alg14 (Gao et al., 2008). The formation of the Agl14/Alg13 complex is mediated by a short C-terminal α-helix of Alg13 in cooperation with the last three amino acids of Alg14 (Gao et al., 2008). In the Alg13 crystal structure this C-terminal helix is associated with itself, as Agl14 is missing. The orientation is predicted to be similar to the linker in MurG and the change is indicated by the black arrow (in Fig 2A) (Gao et al., 2008). The structural model of Agl24 is also predicted to contain a C-terminal helix, which interacts with the N-terminal part of the protein (Fig 2A). We revealed a conserved GGxGGH_14_ motif (Agl24) within the N-terminal sequences of MurG and Alg14 (Fig 1D, S2, and 2B). This motif is located in the cavity between the two different structural domains next to the substrate binding pocket. The crystal structure of MurG (PDB: 3s2u) in complex with UDP-GlcNAc shows the close proximity of the sugar donor to the conserved residues (Fig 2B) (Brown et al., 2013). In proximity to the GGxGGH_14_ motif is the conserved glutamic acid (E_114_) (Fig 1D, S2, and 2B). Interestingly, both these conserved residues are absent in the archaeal MurG-like GT28 family orthologs of pseudomurein-producing Archaea, where aspartic acid (D) and leucine (L) are more frequently found replacing the H_14_ and E_114_ (Fig 1D and S2). We propose that these residues might be required to accommodate the different acceptor molecule.

**Fig 2:**
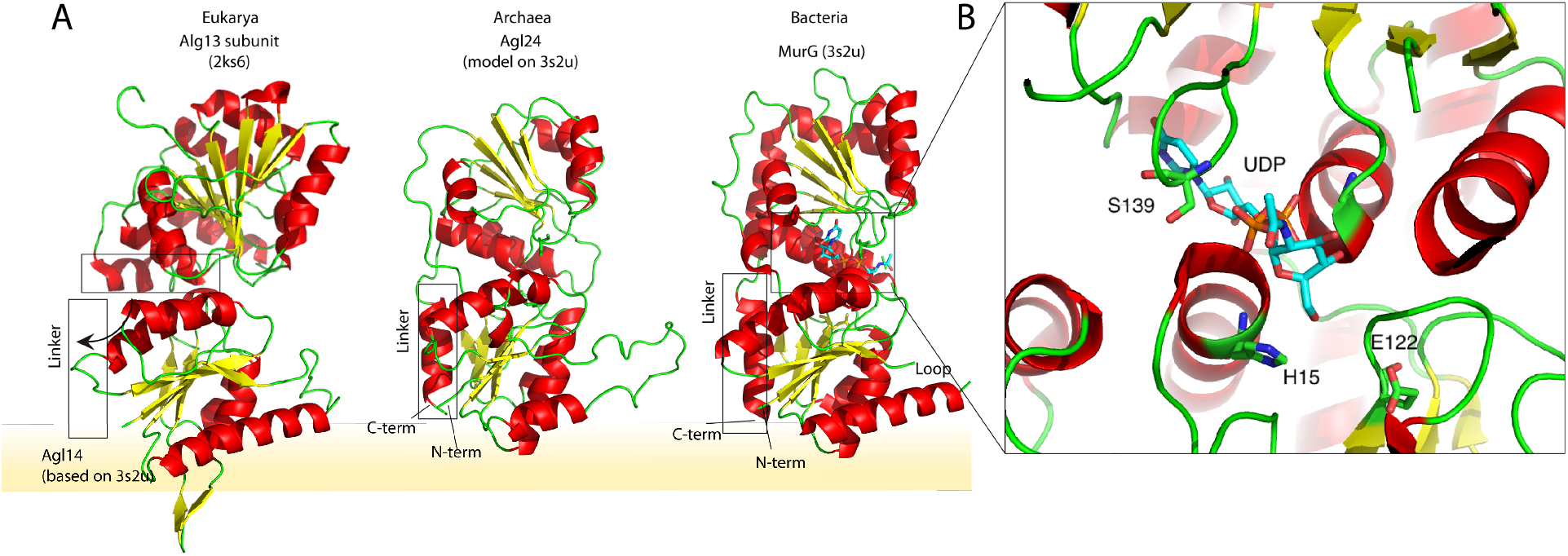
A) Structural comparison of the eukaryal Agl13 (PDB:2ks6) with a structural model of Agl14, the bacterial MurG (PDB: 3s2u), and the structural model of the crenarchaeal Agl24. Structural models were built with SWISS-MODEL (Waterhouse et al., 2018). Detailed results are listed in Supplementary Table S2. The interactions of the N-terminal helices with the membrane are depicted with a yellow background. B) Magnified and 90° rotated view into the active site of MurG (PDB: 3s2u) in complex with UDP-GlcNAc (sticks). The catalytic site is located in the cleft of both domains, with conserved His15 and Glu _(122)_ residues are highlighted in red.

### Agl24 is essential in *S. acidocaldarius*

Due to the overall low sequence similarity of Agl24 to Alg14-13 and MurG, we proposed to demonstrate the predicted function of Agl24 *in vivo* by generating a deletion mutant of *Agl24* in the genome of *S. acidocaldarius*. This deletion mutant should arrest the N-glycosylation process after the generation of Dol-PP-GlcNAc and should contain either non-glycosylated glycoproteins or N-glycosylated proteins containing only a single GlcNAc residue linked to the asparagine in the conserved N-glycosylation motif. The genomic integration by homologous recombination of the plasmid pSVA1312 via either the up- or downstream region of the *agl24* and the selection *pyrEF* genes was confirmed by PCR (Fig S3A). However, we were not successful in generating the marker-less in-frame *agl24* deletion mutant. An alternative approach was conducted, aimed to delete the *agl24* gene by a single homologous recombination step by the integration of the *pyrEF* selection cassette. All attempts to generate this gene disruption mutant failed. Therefore, this strongly suggests that the *agl24* gene is essential in *S. acidocaldarius*, at least under the conditions tested. To exclude that the other GTs, identified in the bioinformatic homology search (Table S1), are involved in the N-glycosylation, marker-less deletion mutants of *saci1907*, *saci1921*, *saci0807, saci1094*, *saci1201*, *saci1821*, and *Saci1249* were successfully generated. Only the deletion of *saci0807* showed a significant effect on the N-glycosylation of the S-layer proteins, which have been characterized to encode the GT Agl16 that transfers the terminal glucose residue to the N-glycan (Meyer et al., 2013).

### Agl24 transfers a single GlcNAc residue onto a GlcNAc acceptor

In agreement with the observed membrane association in *S. acidocaldarius* (Fig S4), Agl24 was also found in the membrane fraction during its purification from *E. coli*. Recombinant Agl24, produced in *E. coli* (Fig S5), was assayed to test the function of Agl24 using the predicted nucleotide sugar donor UDP-GlcNAc and either one of two synthetic acceptor substrates designed to mimic the native lipid-linked acceptor: C_13_H_27_-PP-GlcNAc (acceptor 1) or phenyl-O-C_11_H_22_-PP-GlcNAc (acceptor 2) (Fig 3C). The reaction product was purified and characterized. The MALDI–MS spectra obtained confirmed that Agl24 transfers a single GlcNAc to both acceptor substrates when incubated with UDP-GlcNAc (Fig 5B and D) (Fig 3A). The Agl24 assay revealed that the product peaks are shifted by 203 Da to m/z = 811 [M-1H+2Na]^+^ and m/z = 833 [M-2H+3Na]^+^ (Fig 3B), corresponding to one dehydrated GlcNAc (203 Da). The same shift was observed for acceptor 2 (Fig 3D). We also investigated Agl24 promiscuity towards utilizing UDP-Glucose as the sugar donor, but no mass shift was observed (Fig 3E), indicating that Agl24 uses exclusively UDP-GlcNAc.

**Fig 3:**
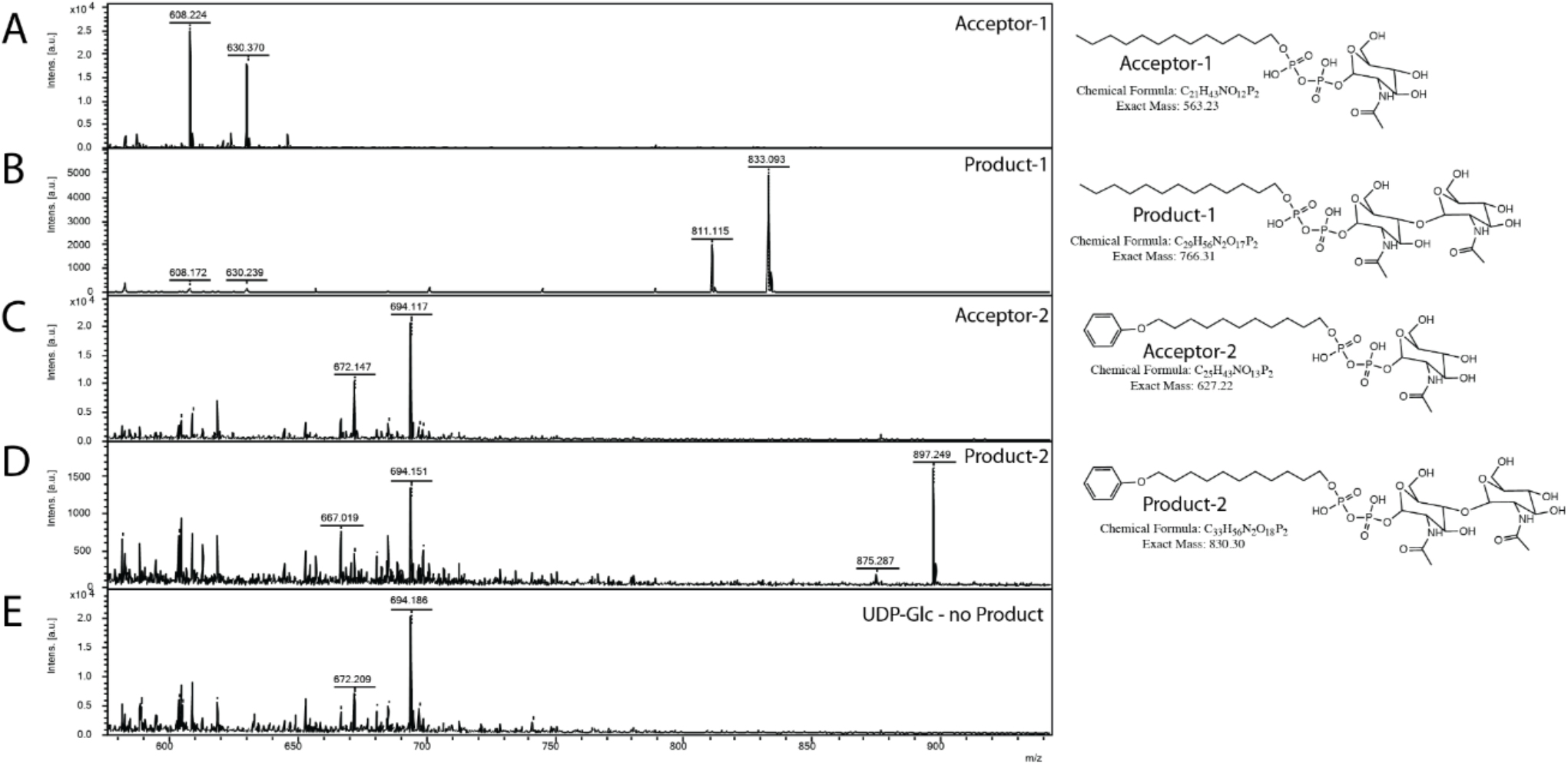
MALDI-MS spectra of the *in vitro* Agl24 reaction assessing the N-acteylglucosamine transferase activity. Spectra obtained from the enzymatic reactions containing Agl24-GFP and (**A**) acceptor-1, **(B)** acceptor-1 and UDP-GlcNAc, and Agl24-GFP, **(C)** acceptor-2, **(D)** acceptor-2, and UDP-GlcNAc, **(E)** acceptor-2 and UDP-Glc. The conversion from acceptor 1 (608 m/z [M-1H+2Na]^+^ and 630 m/z [M-2H+3Na]^+^) or acceptor 2 (672 m/z [M-1H+2Na]^+^ and 694 m/z [M-2H+3Na]^+^) to the product (811 m/z [M-1H+2Na]^+^ and 833 m/z [M-2H+3Na]^+^) or (875 m/z [M-1H+2Na]^+^ and 897 m/z [M-2H+3Na]^+^) was observed only when UDP-GlcNAc was used as nucleotide sugar donor.

### Activity of Agl24

Since *S. acidocaldarius* is a thermophilic microorganism, with an optimal growth temperature of 75°C, the temperature dependency of the Agl24 activity was investigated using our established mass spectrometry assay. The substrates are stable at the conditions tested and MALDI analysis of the negative control lacking the enzyme revealed only the acceptor-1 mass (Fig 4A). At elevated temperatures, the peak intensity from the product increased while the peak intensity of the acceptor molecule was reduced (Fig 4B-F). Highest activity was detected at 70°C, close to the optimal growth temperature of *S. acidocaldarius*. Furthermore, the addition of EDTA did not affect the activity of Agl24, demonstrating that Agl24 is a metal independent GT (Fig S7, 5D). This result is in agreement with the lack of a conserved Asp-X-Asp (DxD) motif within Agl24 sequence, which has been shown to be important to coordinate the metal ions in A-fold GTs, whereas GT-B GTs are metal ion-independent(Lairson et al., 2008, Gloster, 2014).

**Fig 4:**
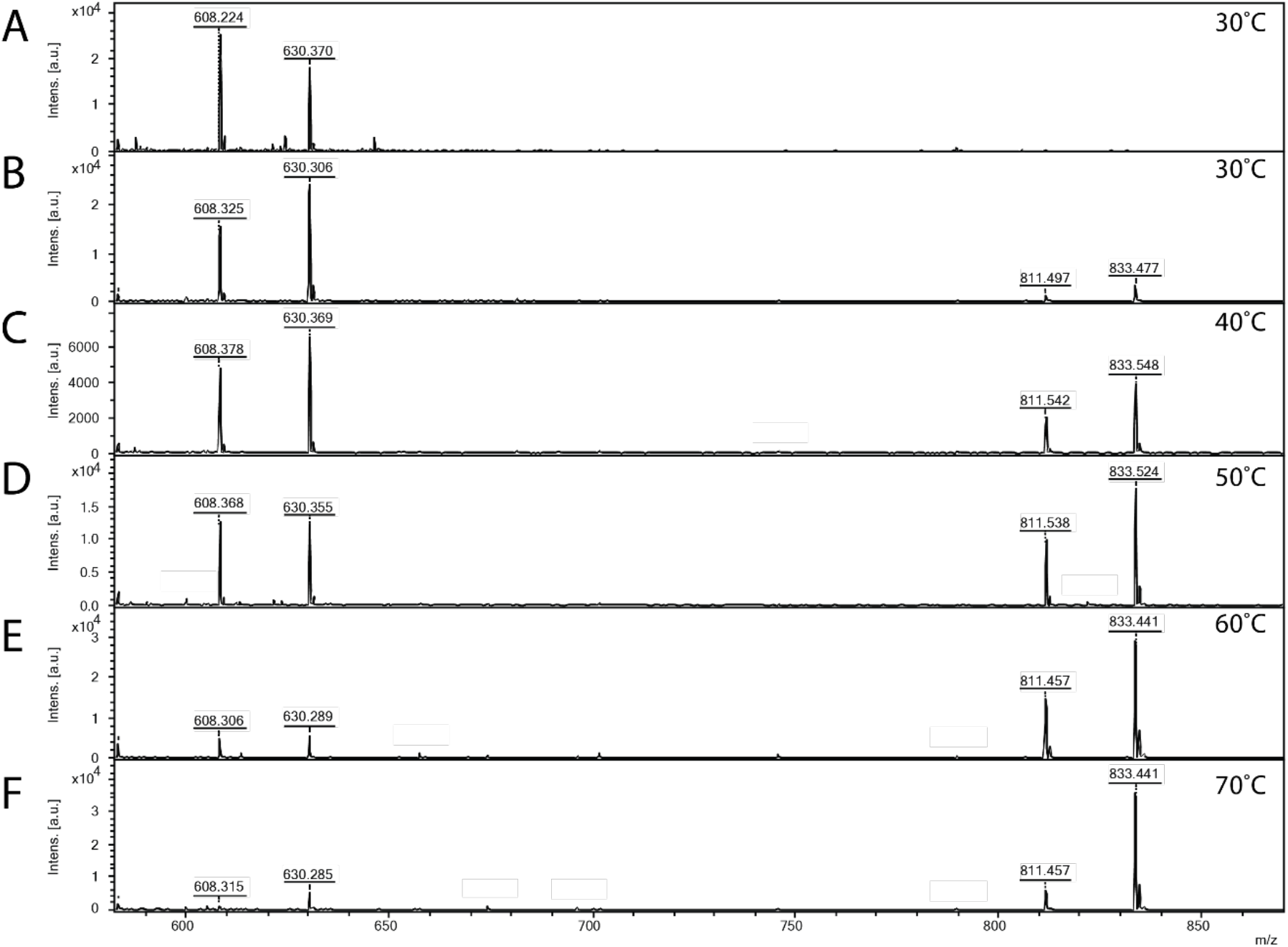
MALDI-MS spectra of the *in vitro* Agl24 reaction at different temperatures. Spectra obtained from the purified enzymatic reaction mix with only the acceptor-1 (**A**), acceptor-1, UDP-GlcNAc, and Agl24-GFP WT at 30°C (**B**), 40°C (**C**), 50°C (D), 60°C (**E**), and 70°C (F). Conversion of the acceptor-1 (608 m/z [M-1H+2Na] and 630 m/z [M-2H+3Na]) towards the product (811 m/z [M-1H+2Na] and 833 m/z [M-2H+3Na]) is significantly dependent on the applied temperature. Above 50°C the depletion of the acceptor towards the gain of product is clearly visible.

### Conserved His14 is essential for Agl24 function

Two conserved amino acid residues were targeted by mutagenesis to investigate their role in the Agl24 enzyme. Alanine substitution of histidine residue H_14_ within the conserved GGH_14_ motif, found in all Alg14 (GSGGH), MurG (GGxGGH), and Agl24 (GGH) homologs, was inactive (Fig 5B). This demonstrated that this highly conserved residue, located next to the nucleotide-binding site, is important for the enzyme function. This GGxGGH motif, and a subsequent second glycine rich motif (GGY), have been proposed to enable MurG to be involved in interaction with the diphosphate group of the lipid acceptor, as these two motifs resemble phosphate-binding loops of nucleotide-binding proteins (Baker et al., 1992, Carugo and Argos, 1997). Alanine substitution of the conserved His residue in MurG resulted in undetectable activity and loss of ability to complement a temperature sensitive MurG mutant (Crouvoisier et al., 2007, Hu et al., 2003). In contrast, the substitution of the second conserved residue E_114_, opposite the nucleotide-binding site, had no effect on the activity of Agl24 (Fig 5C). In MurG this glutamic acid residue is found in a conserved H_124_EQN_127_ motif (*E. coli*), proposed to coordinate the lipid acceptor (Crouvoisier et al., 2007, Hu et al., 2003). The MurG E_125_A variant was able to complement the thermosensitive MurG *E.coli* strain, but the remaining activity was 860-fold lower than that of the wild-type MurG (Crouvoisier et al., 2007).

**Fig 5:**
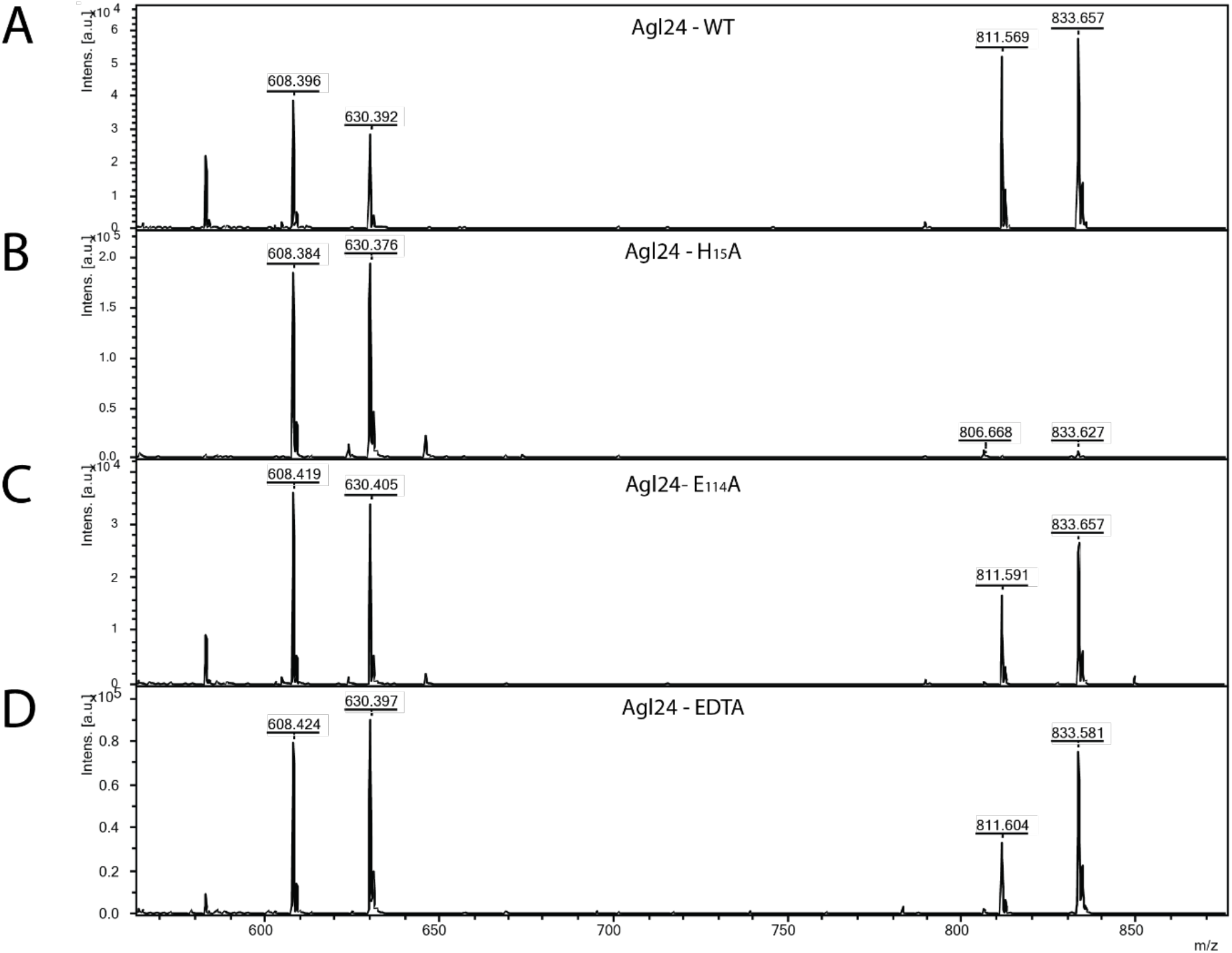
MALDI-MS spectra of the *in vitro* Agl24 reaction, the generated point mutations H15A and E114A and EDTA control. Spectra obtained from the purified enzymatic reaction mix with acceptor-1(608 m/z [M-1H+2Na] and 630 m/z [M-2H+3Na]), UDP-GlcNAc and (**A**) Agl24-GFP WT, (**B**) Agl24-H_15_A-GFP, (**C**) Agl24-E_114_A-GFP, and (**D**) Agl24-GFP WT with addition of 10 mM EDTA. The activity was significantly reduced in the H_15_A mutants, while a similar amount of product (811 m/z [M-1H+2Na] and 833 m/z [M-2H+3Na]) was obtained in the E_114_A mutant.

### Agl24 is an inverting β-1,4-N-acetylglucosamine-transferase

For kinetic analysis of Agl24, a HPLC assay was used to monitor conversion of the mono-GlcNAcylated-PP-lipid acceptor-2 substrate to the bi-GlcNAcylated product (Fig S6). A *Km* value for acceptor-2 could not be determined using this system due to significant substrate inhibition above 0.5 mM, however, a *Km*^(app)^ for UDP-GlcNAc was determined: ^(UDP-GlcNAc)^*Km* = 1.37 ± 0.13 mM, *V*_*max*_ = 32.5 ± 0.9 pmol min^−1^. These results are in agreement with the *Km* values of other GT enzymes, which typically exhibit *K*_m_ affinities for their respective nucleotide sugars in the high micromolar range, reflecting the estimated intracellular concentrations of the nucleotide sugar (Varki et al., 1999). A ^1^H NMR analysis confirmed that the enzymatic product contained two GlcNAc residues with different linkage types based on the presence of two differently-coupled anomeric protons (Fig 6 and S8). One anomeric proton appeared as a doublet of doublets (5.35 ppm, *J*_H1-H2_ = 3.1 Hz, *J*_H1-P_ = 7.2 Hz) typical of an α-linked GlcNAc residue (Table 1). Another anomeric signal appeared as a doublet with a large *J* coupling value (4.49 ppm, *J*_H1-H2_ = 8.5 Hz), indicative of a β-linked GlcNAc residue. The substrate acceptor 2 contains an α-linked GlcNAc; this strongly suggested the terminal GlcNAc introduced by Agl24 was β-linked. Experiments using 1D total correlation spectroscopy (TOCSY) deciphered the detailed proton signals from each of the two sugar rings (Fig S8), and 2D COSY (Fig S9). In addition, HSQC experiments (Fig S10) were used to assign the identity of each proton and carbon signal. The C4 signal of the α-GlcNAc residue is shifted to 79.6 ppm, strongly suggesting that the terminal β-GlcNAc was linked at this position. This was further supported by a relative increase in shift of the H4 proton of the α-GlcNAc residue compared to the un-modified acceptor (Zorzoli et al., 2019). Relative shifts of other protons reported for the acceptor aligned with our experimental data. In conclusion, a combination of ^1^H NMR and 1D and 2D TOCSY, COSY and HSQC experiments confirmed that Agl24 is an inverting β-1,4-GlcNAc transferase.

**Table 1.** ^1^H and ^13^C chemical shifts of the *in vitro* Agl24 reaction product.

| Lipid | 1 | 2 | 3 to 8 | 9 | 10 | 11 | 12, 14 | 13, 15 | 16 |
| --- | --- | --- | --- | --- | --- | --- | --- | --- | --- |
| $^1\text{H}$ NMR | 3.79 | 1.51 | 1.16-1.26 | 1.33 | 1.66 | 3.98 | 6.92 | 7.26 | 6.92 |
| $^{13}\text{C}$ NMR | 66.8 | 30.3 | 28.8 | 28.7 | 28.7 | 68.8 | 115.6 | 130.1 | 121.8 |
| **α-GlcNAc** | **1'** | **2'** | **3'** | **4'** | **5'** | **6'a** | **6'b** |  |  |
| $^1\text{H}$ NMR | 5.35 | 3.896 | 3.8 | 3.58 | 3.83 | 3.73 | 3.57 | | |
| $^{13}\text{C}$ NMR | 94.25 | 53.65 | 70 | 79.65 | 71.69 | 60.15 | | | |
| **β-GlcNAc** | **1''** | **2''** | **3''** | **4''** | **5''** | **6''a** | **6''b** |  |  |
| $^1\text{H}$ NMR | 4.49 | 3.64 | 3.45 | 3.34 | 3.38 | 3.8 | 3.6 | | |
| $^{13}\text{C}$ NMR | 101.55 | 56.14 | 74.01 | 70.38 | 76.27 | 61.15 | | | |

**Fig 6.**
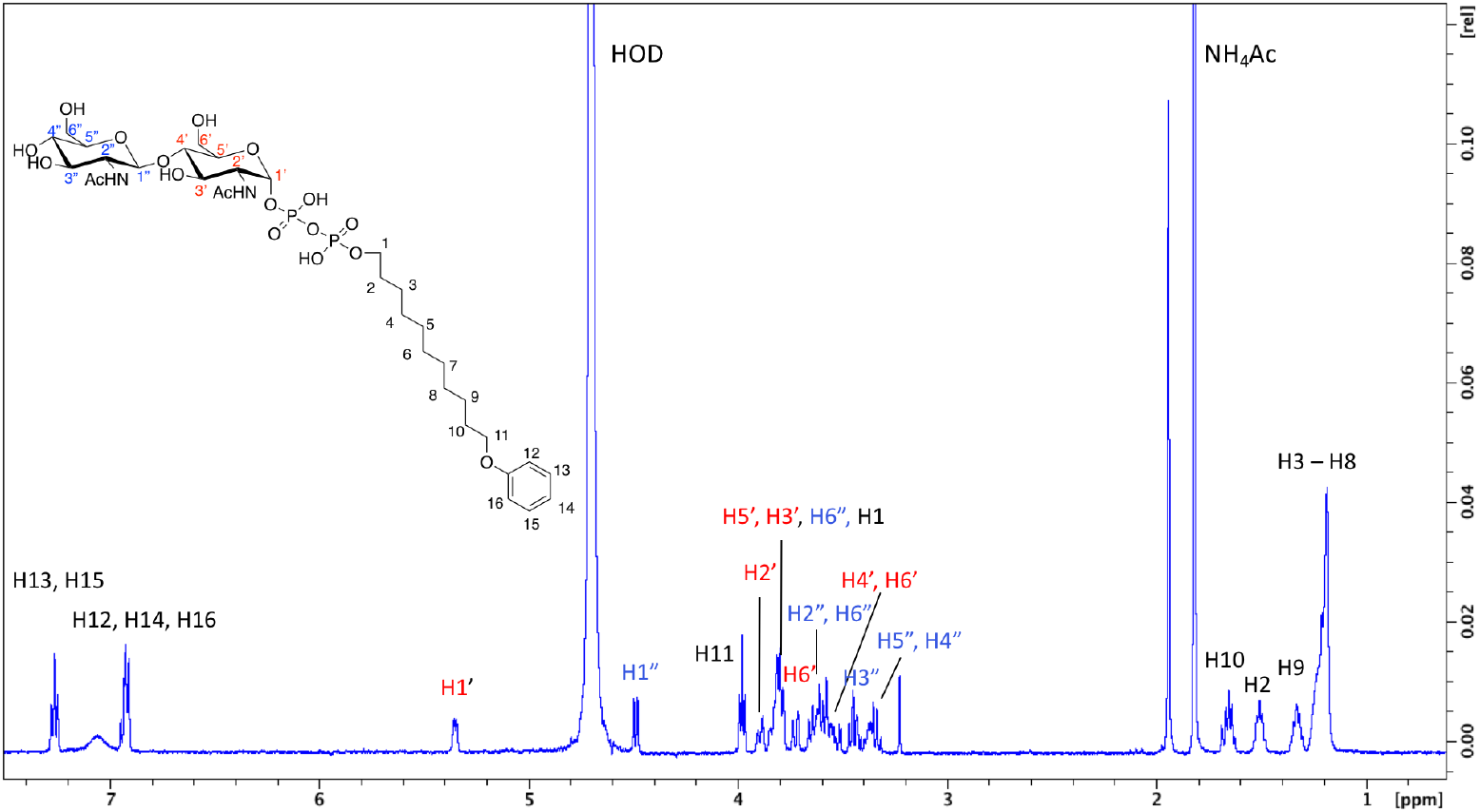
^1^H NMR spectra of purified Agl24 enzymatic product. Spectra were acquired in D2O at 293K on a Bruker AVANCE III HD 500 MHz NMR Spectrometer fitted with a 5-mm QCPI cryoprobe. Chemical shifts are reported with respect to the residual HDO signal at *δ*_H_ 4.70 ppm.

### The eukaryotic GTs Alg14 and Alg13 are closely related to Asgard homologs

In order to analyze the phylogenetic relationship between the eukaryotic N-glycosylation GTs Alg13 and Alg14 with archaeal and bacterial homologs, an extensive phylogenetic analysis was performed.

The eukaryotic Alg13 and Alg14 sequences cluster with homologs from the Asgard phyla Thorarchaeota and Odinarchaeota, as well as with sequences from Verstraetearchaeota, Micrarchaeota, Geothermarchaeota, and an unclassified archaeon (Fig7 A and B). Even though the sister clades of Eykaryotes are Thorarchaeota and Odinarchaeota for Alg13 and Alg14 respectively, the only the latter relationship is strongly supported. No closely-related Alg13- and Alg14-like sequences were found in other Asgard phyla, suggesting that the remaining Asgard, such as Lokiarchaeota and Heimdallarchaeota, could be using a different enzyme for synthesizing N-glycans.

**Fig 7:**
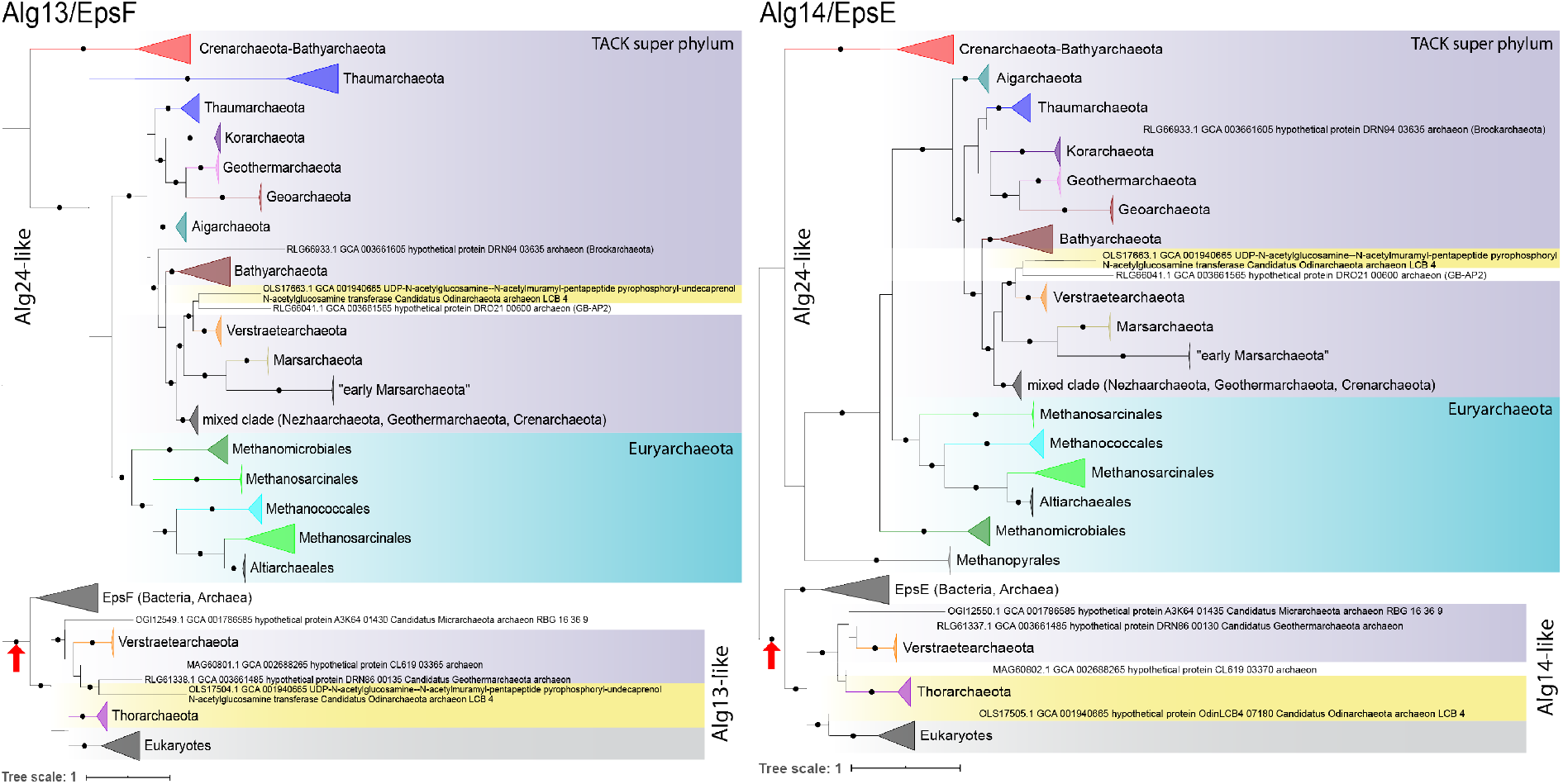
Single gene phylogenies of Alg13/EpsF and Alg14/EpsE. Dots indicate strongly supported branches (ultrafast bootstrap >=95 and aLRT SH-like support >=80). Higher-level classifications are highlighted in specific background colors: purple: archaeal TACK superphylum, turquoise: Euryarchaeota, yellow: Asagradarchaeota, grey: Eukaryotes. Agl24 of *S. acidocaldarius* is found within the red colored clade at the top of the phylogenetic tree. Red arrow: indicates major split event to form the Alg13/EpsF-like and Alg14/EpsE-like enzymes.

Next to the Alg13- or Alg14-like cluster, there is a second distinct cluster of EpsF- or EpsE-like sequences, respectively. The function of EpsF and EpsE has been studied in Lactobacillales, where these enzymes are involved in exopolysaccharide production (De Vuyst et al., 2001, Kolkman et al., 1997). Here, similar to Alg14/13, the combined activity of EpsE and EpsF links either a glucose (Glc) from UDP-Glc (van Kranenburg et al., 1999, Kleerebezem et al., 1999), or a galactose (Gal) from UDP-Gal via a β-1,4 linkage to lipid-linked glucose acceptors (van Kranenburg et al., 1999), creating either a lipid-linked cellobiose or a lactose. The archaeal Eps-like sequences are mainly found in diverse Euryarchaeota, whereas the bacterial sequences are scattered among different phyla, possibly as a result of extensive horizontal gene transfers.

The remaining clades in the Alg13 and Alg14 phylogenies consist of sequences from the archaeal TACK superphylum and some Euryarchaeota, mainly methanogens (Fig 7A and B). The Agl24 identified in this study is found among them, within a Cren- and Bathyarchaeota branch. Close homologs of the Agl24 are found in all presently available genomes of the Crenarchaeota, in the orders Fervidicoccales, Acidilobales, Desulfurococcales and Sulfolobales, with the exception of any homolog in the members of the order Thermoproteales (Fig S11, Sup Data 1).

The remaining phyla of the TACK superphylum (Aig-, Bathy-, Geotherm-, Kor-, Thaum-, Nezha-, Mars-, Brock-, and Verstraetearchaeota) along with some additional Crenarchaeota, including Geoarchaeota, and an Odinarchaeota sequence (Fig 7A and B), form a separate branch containing Agl24-like sequences. Next to this branch is a clade of methanogens and Altiarchaeales. Due to the large number of very distant Alg13/14 homologs with low similarity, resulting in poor alignments, for the phylogenies in Fig 7 we sought to imitate and expand the clades presented previously (Lombard, 2016). Nevertheless, in the homology searches used to expand the previous set (Lombard, 2016), some of the hits formed well-supported branches in preliminary phylogenies (Sup Data 1). By including some of them in expanded Alg13/EpsF and Alg14/EpsE datasets, none of them branch within the Alg13/EpsF-like or Alg14/EpsE-like clades. Instead, they are fused Agl24-like sequences, mainly from Euryarchaeota and various DPANN phyla. Either way, the Cren-/Bathyarchaeota Agl24 clade along with a small Nanohaloarchaeota clade appear to be the closest relatives of the Alg13/EpsF-like and Alg14/EpsE-like proteins (Sup Data 1).

The fact that we are able to trace Agl24-like homologs to the ancestor of TACK and several Euryarchaeota lineages indicates that, despite multiple instances of lateral gene transfer, they are very ancient and diverged from the bacterial MurG at the separation of the two domains, as proposed by Lombard *et al.* (Lombard, 2016). Given that both MurG (Lombard, 2016) and the various Agl24-like homologs are fused genes, a major split event seems to have occurred once at the base of the Alg/Eps-like clade (Fig 8A and B, red arrow), with some independent, more recent, fusions found throughout eukaryotes and Archaea (*e.g.* the Thorarchaeota RLI55694.1). Finally, recovering the monophyly of eukaryotes in our phylogenies corroborates the proposition by Lombard *et al.* (Lombard, 2016) that Alg13 and Alg14 were present in the last eukaryotic common ancestor (LECA). Since the sister clade of eukaryotes is in both cases an Asgard phylum (although not the Heimdallarchaeota), it is possible that these genes were inherited vertically during eukaryogenesis in a two-domain tree of life (Eme et al., 2017).

## Discussion

In this study we identified the first archaeal and thermostable β-1,4-N-acetylglucosaminyltransferase, named <u>a</u>rchaeal <u>g</u>lycosylation enzyme 24 (Agl24), responsible for the synthesis of the lipid-linked chitobiose on which the N-glycan of *S. acidocaldarius* is assembled. The reported essentiality of Agl24 is in agreement with the indispensable properties of the OST AglB and AglH, catalyzing either the last or first step of the N-glycosylation process in *S. acidocaldarius* (Meyer and Albers, 2014, Meyer et al., 2017). The essentiality of the N-glycosylation has also been confirmed by a transposon study in *Sulfolobus islandicus*, a close relative to *S. acidocaldarius*, in which all homologous genes, including *agl24*, lack any transposon insertions (Zhang et al., 2018). However, the fundamental nature of archaeal N-glycosylation cannot be generalized since deletion mutants of the oligosaccharyltransferase AglB can be obtained in Euryarchaeota, *e.g.,* in *Methanococcus maripaludis* and *Haloferax volcanii* (Abu-Qarn et al., 2007, Chaban et al., 2006). The requirement for N-glycosylation might rely on many factors including the N-glycan composition and structure as well as the modification frequency on proteins; this frequency is reported to be higher in thermophilic, compared to mesophilic, Archaea (Meyer and Albers, 2013, Jarrell et al., 2014).

The comparison of Agl24 with eukaryotic Alg14/13 and the bacterial MurG enzymes revealed structural and amino acid similarities. Although the sequence identity of Agl24 with the eukaryotic and bacterial enzymes is extremely low (~16%), the structural prediction of Agl24 indicated a MurG–like GT-B fold. Interestingly, the amino acids of the nucleotide sugar donor binding site of MurG (Hu et al., 2003) are not conserved in Agl24 and Alg14, although all enzymes use UDP-GlcNAc as the sugar donor (Fig S2, boxed). Strikingly, a conserved GGH_14_ motif within an N-terminal loop of MurG, Alg14-13, and Agl24 (Fig 1 and S2), which protrudes into the cavity formed by the two structural folds, could be identified. In Bacteria this G_13_GTGGH_18_ loop is extended by a second G_103_GY_105_ loop. As both loops are suggested to be reminiscent of the phosphate-binding loops of nucleotide-binding proteins, their involvement in the binding of the diphosphate group of the acceptor lipid has been proposed (Crouvoisier et al., 2007). In agreement with our results, the alanine substitutions of the invariant bacterial H_14_ led to an extremely low or undetectable activity of MurG as well as the loss of ability to complement a temperature sensitive MurG mutant (Crouvoisier et al., 2007, Hu et al., 2003). However, up to now, no structure of MurG or Agl14-13 with the bound lipid acceptor molecule has been published and the interaction of the imidazole ring of the histidine residue with the acceptor molecule remained to be shown. Our alignment also reveals a conserved glutamic acid in the bacterial, eukaryotic, and archaeal sequences. In bacteria, this glutamic acid is found in a conserved H_124_EQN_127_ loop (*E. coli* MurG), which has been proposed to form a key part of the acceptor-binding site (Hu et al., 2003). In agreement with our study, an E_125_A substitution significantly affected the function of MurG, although a marked decrease in lipid acceptor binding (1760-fold) and nucleotide substrate binding (210-fold) was observed (Hu et al., 2003, Crouvoisier et al., 2007).

Although lacking a TMD, Agl24 associates with the membrane, indicating interaction with membrane enzymes, similar to the interaction of MurG or Alg13/14 with their membrane interaction partners MraY and Alg7 respectively. Based on the structural similarities of these complexes, a shared common evolutionary origin of the first enzymes of eukaryotic N-glycosylation and the bacterial peptidoglycan biosynthesis has been hypothesized (Bugg and Brandish, 1994, Burda and Aebi, 1999, Bouhss et al., 2008). As *S. acidocaldarius* possesses a homolog of Alg7, termed AlgH (Meyer et al., 2017), and a bioinformatics study has indicated that Alg7 might have emerged out of the Crenarchaeota (Lombard, 2016), AglH presents the most likely candidate for the interaction partner of Agl24.

A phylogenetic analysis shed light onto the distribution of the identified Agl24 in Archaea, and determined its relationship with eukaryotic and bacterial homologs. This revealed a very ancient and broad distribution of Agl24-like sequences in Archaea. Importantly, close homologs of Agl24 are only found within the Crenarchaeota and Bathyarchaeota. Indeed, all Crenarchaeota, with the exception of the order Thermoproteales, possess an Agl24 homolog, strongly suggesting that these Crenarchaeota contain a chitobiose N-glycan core structure. In contrast, the N-glycans of members of the Thermoproteales would be assembled using a different enzyme. Indeed, the N-glycan of *Pyrobaculum caldifontis*, belonging to the order Thermoproteales, differs from the N-glycan in Sulfolobales (Fujinami et al., 2017). Here, the N-glycan core is composed of two di-N-acetylated β-1,4-linked GlcNAc residues with the second sugar carboxylated at C6 (GlcA(NAc)_2_-β-1,4-(Glc(NAc)_2_. A di-N-acetylated glucuronic acid sugar is also found in the N-glycan from *Methanococcus voltae,* which is transferred by the GT AglC onto the Dol-P linked GlcNAc (Chaban et al., 2009). Thus, a homolog of AglC is likely to be involved in the N-glycan biosynthesis in *P. caldifontis*.

Our phylogenetic analysis revealed that the closest related enzymes of the eukaryotic Alg14 and Alg13 are found within the Asgard superphylum. Since the discovery of the Archaea as the third domain of life next to Bacteria and Eukarya (Woese and Fox, 1977), the comparison of archaeal molecular processes has gradually revealed a strong resemblance to those found in eukaryotes (Zillig et al., 1989, Huet et al., 1983). Furthermore, recent phylogenetic analyses indicated that eukaryotes originated close to the TACK superphylum (Thaumarchaeota, Aigarchaeota, Crenarchaeota, and Korarchaeota) within the Asgard superphylum (Spang et al., 2015, Zaremba-Niedzwiedzka et al., 2017, Petitjean et al., 2015, Koonin, 2015). The current hypothesis for the formation of the first eukaryotic cell proposes a membrane protrusion of an ancestral archaeon that engulfed a bacterium to increase the interspecies interaction surface (Baum and Baum, 2014, Imachi et al., 2020). The eukaryotic N-glycosylation would therefore be inherited from this ancient archaeal ancestor, which is in agreement with the high conservation of the initial N-glycosylation biosynthesis steps in all eukaryotes (Helenius and Aebi, 2004, Samuelson et al., 2005, Schwarz and Aebi, 2011). In addition, the AglB in members of the Asgard and TACK superphyla shows considerable similarity to its eukaryotic homologue Stt3, and is clearly distinct from the AglB of the DPANN superphylum and the Euryarchaeota (Nikolayev et al., 2020). Most eukaryotic Stt3 contain a double sequon motif for N-glycosylation near the substrate-binding site, which is hypothesized to contribute to the LLO (Shrimal and Gilmore, 2019, Wild et al., 2018). Interestingly, this double sequon is also present in the AglB of the members of the Asgard and TACK superphyla but absent in the AglB of Euryarchaeota (Shrimal and Gilmore, 2019). In addition, homologs of non-catalytic subunits of the eukaryotic Stt3 complex, *e.g.,* ribophorin, Ost3/6, Ost5, and Wbp1, have been detected in some Arsgardarchaeota (Zaremba-Niedzwiedzka et al., 2017). However, their specific function in the N-glycosylation has to be shown, especially as ribophorin is only found in higher mammals (Wilson and High, 2007) and therefore unlikely to exist in an early eukaryotes. Nevertheless, the assembly of the N-glycan linkage in the TACK superphylum is conducted via a pyrophosphate linkage to the Dol, identical to that in Eukarya. This is in contrast to Euryarchaeota, which use a mono-phosphate linkage. A summary of the similarities and differences of the archaeal N-glycosylation to Eukarya and Bacteria are given in Table S4. All these observations strengthen the hypothesis that the eukaryotic N-glycosylation has emerged from an ancient archaeon.

In the future, we believe that the detailed characterization of the N-glycosylation process in members of the TACK and Asgard superphyla will lead to the elucidation of further molecular similarities and unique properties to the eukaryotic N-glycosylation process and will provide further support for the origin of the eukaryotic N-glycosylation.

## Materials and methods

### Strains and growth conditions

The strain *Sulfolobus acidocaldarius* MW001 (Δ*pyrE*) (Wagner et al., 2012) and all derived modified strains (Table S3) were grown in Brock medium at 75°C, pH 3, adjusted using sulfuric acid. The medium was supplemented with 0.1% w/v NZ-amine and 0.1% w/v dextrin as carbon and energy source (Brock et al., 1972). Selection gelrite (0.6%) plates were supplemented with the same nutrients with the addition of 10 mM MgCl_2_ and 3 mM CaCl_2_. For second selection plates 10 mg ml^−1^ uracil and 100 mg ml^−1^ 5-fluoroorotic acid (5-FOA) were added. For the growth of the uracil auxotrophic mutants, 10 mg ml^−1^ uracil was added to the medium. Cell growth was monitored by measuring the optical density at 600 nm. Protein expression in *S. acidocaldarius* was conducted in medium supplemented with 0.1% w/v NZ-amine and 0.1% w/v L-arabinose as carbon and energy source. All *E. coli* strains DH5α, BL21, or ER1821 were grown in LB media at 37°C in a shaking incubator at 200 rpm. According to the antibiotic resistance in the transformed vector(s), media were supplemented with the antibiotics carbenicillin (amp) at 100 μg ml^−1^ and/or kanamycin (kan) at 50 μg ml^−1^.

### Construction of deletion plasmids

The predicted function of *Agl24* was verified via the generation of the lipid-linked chitobiose core of the *N*-glycan. Therefore, a marker-less deletion mutant of this gene was constructed in the *S. acidocaldarius* MW001, as previously described (Wagner et al., 2012). Briefly, the strain MW001, auxotrophic for uracil biosynthesis, was transformed with the plasmid pSVA1312. Therefore, two ~1000 bp DNA fragments, one from the upstream and one from the downstream regions of *Agl24* (*saci1262*) gene, were PCR amplified. Restriction sites *Apa*I and *Bam*HI were introduced at the 5’ ends of the upstream forward primer (4168) and of the downstream reverse primer (4165), respectively. The upstream reverse primer (4163) and the downstream forward primer (4164) were each designed to incorporate 15 bp of the reverse complement strand of the other primer, resulting in a 30 bp overlap stretch. The overlapping PCR fragments were purified, digested with *Apa*I and *Bam*HI, and ligated in the digested plasmid pSVA407, containing *pyrEF* (Wagner et al., 2012).

### Generation of a linear *Agl24*_up_-*pyrEF*-*Agl24*_down_ fragment for the direct *Agl24*::*pyr* EF replacement

To further underline the essential role of *Agl24* in *S. acidocaldarius*, a disruption of the *Agl24* gene by direct homologous integration of the *pryEF* cassette was performed. For this approach, 387 bp of the *Agl24* upstream region, the full 1525 bp of the *pyrEF* cassette, and 1011 bp of the *Agl24* downstream region were PCR amplified using the primers: 4168+6336, 4115+4116, and 6337+6338, respectively. At the 5’ ends of the upstream forward primer (4168) and of the downstream reverse primer (6338) restriction sites *Apa*I and *Bam*HI were introduced, respectively. The *pyrEF* forward primer (4115) was designed to incorporate 40 bp of the upstream reverse primer (6336) resulting in a 40 bp overlapping stretch. The *pyrEF* reverse primer (4116) and the downstream forward primer (6337) weres designed to incorporate the reverse complement strand of the other primer, resulting in a 46 bp overlapping stretch. The upstream, *pyrEF*, and downstream fragments were fused by an overlapping PCR, using the 3ʹ ends of each fragment as primers. The 2904 bp overlap PCR fragment gained, was amplified using the outer primers (4168 and 6338), digested with *Apa*I and BamHI and ligated into the p407 vector predigested with the same restriction enzymes. The resulting plasmid, pSVA3338 (*Agl24*_up_-*pyrEF*-*Agl24*_down_), was transformed into *E. coli* DH5α and selected on LB-plates containing 50 mg ml^−1^ ampicillin. The accuracy of the plasmid was verified by sequencing. Before transformation in *S. acidocaldarius,* the plasmid was digested with *Apa*I and *Bam*HI to create the linear *Agl24*_up_-*pyrEF*-*Agl24*_down_ fragment.

### Generation of Agl24 Expression plasmids for the production of Agl24 in *S. acidocaldarius* and *E. coli*

Cellular localization studies were carried out using N- and C-terminal fusion proteins of Agl24. The *Agl24* gene was amplified from genomic DNA of *S. acidocaldarius* introducing *NcoI* and *Pst*I restriction sites at the 5ʹends of the primer pair 4176 and 4177, respectively. The PCR fragment was cloned into pSVA1481, yielding vector pSVA1336 containing an inducible arabinose promoter and *Agl24*-Strep-His_10_. The plasmid was digested with *Nco*I and *Eag*I and the insert *Agl24*-Strep-His_10_ was ligated into pSVA1481 containing an inducible maltose promoter, yielding pSVA1337. The N-terminal Strep-His_10_-TEV-Agl24 was cloned using the primer pair 4180 and 4181, which incorporated the restriction sites *NcoI* and *NotI*, respectively. The Insert was ligated into pSVA2301 containing an inducible maltose promoter yielding pSVA1339.

For heterologous expression of Agl24 in *E. coli,* the plasmid pHD0499 was generated. The GFP-His_8_-tagged *Agl24* gene was constructed by amplifying the full-length *Agl24* gene from *S. acidocaldarius* genomic DNA with the primer pair A596 and A597. The PCR fragment was cloned in-frame into the vector pWaldo (Waldo et al., 1999) using *Xho*I and *Kpn*I restriction sites, generating a C-terminal fusion with a TEV cleavage site, GFP, and His_8_ tag. For the generation of Agl24 mutants of the conserved mutants H_15_ and E_114_, a quick-change mutagenesis PCR was applied using the primers: A597 and A684 or A685 and A686, respectively.

### Transformation and selection of the deletion mutant in *S. acidocaldarius*

Generation of competent cells was performed based on the protocol of Kurosawa and Grogan (Kurosawa and Grogan, 2005). Briefly, *S. acidocaldarius* strain MW001 was grown to an OD_600_ between 0.1 and 0.3 in Brock medium supplemented with 0.1% w/v NZ-amine and 0.1% dextrin. Cooled cells were harvested by centrifugation (2000 x g at 4°C for 20 min). The cell pellet was washed three times successively in 50 ml, 10 ml and 1 ml of ice-cold 20 mM sucrose (dissolved in demineralized water) after mild centrifugation steps (2000 x g at 4°C for 20 min). The final cell pellet was resuspended in 20 mM sucrose at an OD_600_ of 10.0 and stored in 50 μl aliquots at −80°C. 400–600 ng of methylated pSVA1312 or the linearized *Agl24*_up_-*pryEF*-*Agl24*_down_ fragment, was added to a 50 μl aliquot of competent MW001 cells and incubated for 5 min on ice before transformation in a 1 mm gap electroporation cuvette at 1250 V, 1000 Ω, 25 mF using a Bio-Rad gene pulser II (Bio-Rad, X, USA). Directly after transformation 50 μl of a 2x concentrated recovery solution (1% sucrose, 20 mM b-alanine, 20 mM malate buffer pH 4.5, 10 mM MgSO_4_) was added to the sample and incubated at 75°C for 30 min under mild shaking conditions (150 rpm). Before plating, the sample was mixed with 100 μl of heated 2x concentrated recovery solution and twice 100 μl were spread onto gelrite plates containing Brock medium supplemented with 0.1% NZ-amine and 0.1% dextrin. After incubation for 5–7 days at 75°C, large brownish colonies were used to inoculate 50 ml of Brock medium containing 0.1% NZ-amine and 0.1% dextrin, which were incubated for 3 days at 78°C. Cultures confirmed by PCR to contain the genomically integrated plasmid were grown in Brock medium supplemented with 0.1% NZ-amine and 0.1% dextrin to an OD of 0.4. Aliquots of 40 μl were spread on second selection plates, supplemented with 0.1% NZ-amine and 0.1% dextrin and 10 mg ml^−1^ uracil, were incubated for 5–7 days at 78°C. Newly formed colonies were streaked on fresh second selection plates to ensure single colony selection before each colony was screened by PCR for the genomic absence, presence, or modification, of the *Agl24* gene.

### Expression of Agl24 in *S. acidocaldarius*

Plasmid transformation into *S. acidocaldarius* cells was performed as described above for the deletion mutants. Cells were spread onto gelrite plates containing Brock medium supplemented with 0.1% NZ-amine and 0.1% dextrin. After incubation for 5–7 days at 75°C large brownish colonies were used to inoculate 50 ml of Brock medium containing 0.1% NZ-amine and 0.1% dextrin and incubated for 3 days of 78°C. Presence of the expression plasmid was confirmed by PCR. For induction of expression the strains were grown either in Brock medium supplemented with either 0.1% NZ-amine and 0.1% dextrin (MAL_promotor_) or with 0.1% NZ-amine and 0.1% L-arabinose (ARA_promotor_) to an OD_600_ of 1.0.

### Expression and purification of recombinant Agl24 protein from *E. coli*

For heterologous Agl24 protein expression, 6 x 1 Liter LB medium was inoculated with 10 ml from an overnight BL21 culture previously transformed with pHD0499. Cells were grown at 16°C overnight in auto-induction medium (Studier, 2014) containing 30 μg ml^−1^ kanamycin. The cells were harvested by centrifugation at 4,200 x g for 25 minutes and used to isolate the membrane fractions. All subsequent purification steps were carried out at 4°C. Cells were fractionated by passing four times through an Avestin C3 High Pressure Homogeniser (Biopharma, UK), followed by a 20 min low speed spin at 4,000 x g. The resulting supernatant was centrifuged at 200,000 x g for 2 h to obtain the membrane fraction and 2–3 g of membranes were routinely used for isolation of Agl24-GFP-His_8_ proteins. Samples were solubilized in 18 ml Buffer 1 (500 mM NaCl, 10 mM Na_2_HPO_4_, 1.8 mM KH_2_PO_4_ 2.7 mM KCl, pH 7.4, 20 mM imidazole, 4 mM TCEP) with the addition of 1% n-dodecyl-β-maltoside (DDM) for 2 h at 4°C. The sample was two-fold diluted with Buffer 1 and centrifuged at 200,000 x g for 2 h. The supernatant was loaded onto a Ni-Sepharose 6 Fast Flow (GE Healthcare) column with 1 ml of prewashed Ni-NTA-beads. The column was washed with 20 ml of wash buffer (500 mM NaCl, 10 mM Na_2_HPO_4_, 1.8 mM KH_2_PO_4_ 2.7 mM KCl, pH 7.4, 20 mM imidazole, 0.4 mM TCEP, 0.03% DDM) and eluted with 5 x 1ml elution buffer (500 mM NaCl, 10 mM Na_2_HPO_4_, 1.8 mM KH_2_PO4 2.7 mM KCl, pH 7.4, 250 mM imidazole, 0.4 mM TCEP, 0.03% DDM). Elution fractions were combined and imidazole removed using a HiPrep 26/10 desalting column (GE Healthcare) equilibrated with Buffer (1x PBS, 0.03% DDM, 0.4 mM TCEP). Protein concentration was determined using a Bradford reaction (Bio-Rad) and purity was confirmed by SDS– PAGE analysis. The concentrated fractions were separated by SDS–PAGE and stained with Coomassie blue. Protein identity was confirmed by tryptic peptide mass fingerprinting, and the level of purity and molecular weight of the recombinant protein was determined by matrix-assisted laser desorption/ionization time-of-flight mass spectrometry (MALDI–TOF). The analysis was provided by the University of Dundee ‘Fingerprints’ Proteomics Facility.

### Agl24 activity

Activity was measured in a 100 μl reaction volume containing 1 mM UDP-D-GlcNAc, 1 mM acceptor molecule, 5 mM MgCl_2_, and 5 μg of purified Agl24 in TBS Buffer (150 mM NaCl, 50 mM Tris-HCl, pH 7.6); reaction was performed in a PCR cycler at 60°C for 12 h.

### MS analysis

MALDI–TOF was used to analyze the acceptors and products of the Agl24 *in vitro* assay. The 100ul reaction samples were purified over 100 mg Sep-Pak C18 cartridges (Waters) pre-equilibrated with 5% EtOH. The bound samples were washed with 800 μl of H_2_O and 800 μl of 15% EtOH, eluted in two fractions with (a) 800 μl of 30% EtOH and (b) 800 μl of 60% EtOH. The two elution fractions were dried in a SpeedVac vacuum concentrator and resuspended in 20 μl of 50% MeOH. A 1 μl sample was mixed with 1 μl of 2,5-dihydroxybenzoic acid matrix (15 mg ml^−1^ in 30:70 acetonitrile, 0.1% TFA), and 1 μl was added to the MALDI grid. Samples were analyzed by MALDI in an Autoflex Speed mass spectrometer set up in reflection positive ion mode (Bruker, Germany).

### HPLC analysis

With the exception of kinetics reactions, which are detailed below, Agl24 reactions were analyzed using a HPLC assay with, typically, 50 μl samples containing 2.5 mM UDP-GlcNAc, 1.5 mM lipid acceptor, and 1.8 μg of purified Agl24 in a TBS buffer containing 5 mM MgCl_2_ (150 mM NaCl, 50 mM Tris-HCl, pH 7.5). Reactions were left at 60°C (or alternative temperatures) for the desired time period and quenched with one equivalent of acetonitrile to precipitate Agl24. Following filtration to remove precipitate, reactions were injected onto an XBridge BEH Amide OBD Prep column (130 Å, 5 μM, 10 x 250 mm) at a flow rate of 4 ml min^−1^ using a Dionex UltiMate 3000 system (Thermo Scientific) fitted with a UV detector optimized to detect the *O*-phenyl functional group of the acceptor molecule at 270 nm. Each run was 35 minutes using running Buffer A (95% acetonitrile, 10 mM ammonium acetate, pH 8) and Buffer B (50% acetonitrile, 10 mM ammonium acetate, pH 8). A linear gradient from 20–80% buffer B was performed over 20 min, followed by an immediate drop back to 20% buffer B for the remaining 15 minutes of the run to re-equilibrate the column to starting conditions. The Agl24 reaction substrates and products typically eluted 8–11 minutes into a run. For kinetic analyses, assays were performed in triplicate (with the exception of 1 mM conc. which was only performed in duplicate) and altered to contain a fixed concentration of lipid-acceptor (0.5 mM) whilst varying the concentration of UDP-GlcNAc (0.5, 0.75, 1, 1.5, 3, 6, 9, 12 mM). Reactions were run for 10 hours before being quenched and, after completion, areas of substrate and product peaks were calculated to determine the reaction conversion. Conversion over 10 hours was converted to pmol per minute, and the resulting data were analyzed using GraphPad Prism v8 to generate a Michaelis–Menten curve and resulting kinetic data.

### NMR Analysis

The HPLC-purified Agl24 products (0.5mg–2mg) were dried using a Christ RVC 2-25 speed vacuum fitted with a Christ CT 02-50 cold-trap to remove excess acetonitrile, then freeze dried (Alpha 1-2 LDplus, Christ) to remove residual water. Products were subsequently dissolved in 600 μl of D_2_O and NMR spectra were recorded at 293K. The spectra were acquired on a Bruker AVANCE III HD 500 MHz NMR Spectrometer equipped with a 5 mm QCPI cryoprobe. For 1D TOCSY experiments, H1’ was irradiated at 5.35 ppm, H2’ was irradiated at 3.90 ppm, and H1” was irradiated at 4.49 ppm (Fig S8). A combination of ^1^H ^1^H COSY, 1D TOCSY, and ^1^H ^13^C HSQC experiments were used to fully assign the ^1^H and ^13^C signals for the Agl24 reaction product. Full ^1^H and ^13^C chemical shift assignments can be found in Table 1 and are recorded with respect to the residual HDO signal at 4.7 ppm.

### Phylogenetic analysis

The eukaryotic Alg13-Alg14 and the bacterial EpsF-EpsE sequences were taken from the datasets in (Lombard, 2016), since our own databanks (25118 bacterial and 1611 eukaryotic genomes) contained too many and too divergent hits that could possibly render downstream analyses unfeasible. For Archaea, HMM searches were impossible, since they recovered too many and too divergent possible homologs to handle, while DIAMOND searches were not sensitive enough. Ultimately, we searched for homologs with DIAMOND (--more-sensitive −k 0 & default e-value cutoff) (Buchfink et al., 2015) using multiple seeds from (Lombard, 2016) to cover the different clades of archaeal homologs: 1) the *S. acidocaldarius* Agl24 from this study; 2) the Alg13/EpsF-like and Alg14/EpsE-like homologs from *Methanothrix soehngenii* (NCBI: AEB67939.1 and AEB67938.1 respectively); 3) AIC14646.1 from *Nitrososphaera viennensis* (TACK clade); 4) ABX11947.1 from *Nitrosopumilus maritimus* (additional Alg13/EpsF-like Thaumarchaeota clade); 5) AAB99267.1 from *Methanocaldococcus jannaschii* (methanogen clade). For the first two, we repeated the homology search with the same parameters using all the hits from the previous round as seeds to ensure we did not miss any sequences from our primary clades of interest. The hits from each search were aligned with MAFFT L-INS-I v7.453 (Katoh and Standley, 2013) and preliminary phylogenies were reconstructed with IQ-TREE 2.0.5 (Minh et al., 2020) (-m TEST −alrt 1000). From these phylogenies we isolated the sequences forming well-supported branches corresponding to the ones in (Lombard, 2016), albeit with increased diversity plus some additional clades of potential interest. The individual datasets (archaeal, bacterial, eukaryotic) were then fused into single Alg13/EpsF-like and Alg14/EpsE-like datasets that were aligned as above and trimmed with ClipKIT (kpigappy) (Steenwyk et al., 2020). Then phylogenies were inferred with IQ-TREE 2.0.5 (Minh et al., 2020) under the model selected by ModelFinder (Kalyaanamoorthy et al., 2017), and ultrafast bootstrap (Hoang et al., 2018), aLRT SH-like (Guindon et al., 2010), and approximate Bayesian (Anisimova et al., 2011) branch support tests (-m MFP −bb 1000 −alrt 1000 −abayes). The trees were visualized in iTol (Letunic and Bork, 2019).

## Abbreviation

Agl: archaeal glycosylation enzyme
Alg: Asparagine linked glycosylation
CAZy: Carbohydrate-Active enZymes database
Dol: dolichol
Dol-P: dolichol-phosphate
DPANN: archaeal superphylum containing Diapherotrites, Parvarchaeota, Aenigmarchaeota, Nanoarchaeota, Nanohaloarchaeota, Woesearchaeota and Pacearchaeota
Glc: glucose
GlcA: glucuronic acid
GlcNAc: N-acetylglucosamine
GFP: green fluorescent protein
GT: glycosyltransferase
LLO: lipid-linked oligosaccharide
MALDI-MS: matrix-assisted laser desorption ionization mass spectrometry
OST: oligosaccharyltransferase
SDS-PAGE: sodium dodecyl sulfate–polyacrylamide gel electrophoresis
TACK: archaeal superphylum containing Thaumarchaeota, Aigarchaeota, Crenarchaeota and Korarchaeota
TMD: transmembrane domain
UDP: uridine diphosphate
Und: undecaprenol

## Author contributions

B. H. M., S. V. A., and H. C. D. conceptualization; B. H. M. data curation, initial BLAST searches, deletion analysis, protein alignments, protein expression, protein purification, alanine substitutions, activity analyses, MALDI analyses, visualization; B. A. W. HPLC activity analyses and NMR analyses; P. S. A. phylogenetic analyses, B. H. M‥, B. A. W., P. S. A. and H. C. D. investigation; B. H. M. and H. C. D. validation; B. H. M. and H. C. D. methodology; B. H. M. writing-original draft; B. H. M., P. S. A., B. A. W., S. V. A., and H. C. D. writing-review and editing; H. C. D. supervision; H. C. D. and S. V. A. funding acquisition

## Acknowledgments

We thank Vladimir I. Torgov, Vladimir V. Veselovsky, and Leonid L. Danilov from the N. D. Zelinsky Institute of Organic Chemistry, Russian Academy of Sciences, Moscow, Russia for sharing the synthetic acceptor molecules. We thank Prof. Alexander J. Probst from the University Duisburg-Essen for fruitful discussions and Mr. Tom Snelling for assay assistance.

