## Supplementary Data (Tables S1-S4 and Figures S1-S11) for "N-glycan chitobiose core biosynthesis by Agl24 strengthens the hypothesis of an archaeal origin of the eukaryal N-glycosylation"

**Table S1 Results of the Domain Enhanced Lookup Time Accelerated BLAST (https://blast.ncbi.nlm.nih.gov/Blast.cgi).** DELTA BLAST was performed to identify homologs of MurG (WP_063074721.1), Alg13 (NP_011468.1), Alg14 (NP_009626.1), and Alg14-13 fusion in *Sulfolobus acidocaldarius*.

| Description | Max score | Total score | Query cover | E value | Per. Ident | Accession | Saci gene | KO | Annotation |
|---|---|---|---|---|---|---|---|---|---|
| **MurG** WP_063074721.1 undecaprenyldiphospho-muramoylpentapeptide beta-N-acetylglucosaminyltransferase [*Escherichia coli*] | | | | | | | | | |
| conserved Archaeal protein [Sulfolobus acidocaldarius DSM 639] | 77.0 | 77.0 | 76% | 6e-17 | 14.52% | AAY81210.1 | *saci_1907* | MW061 BM-A045-B027 | RFaB GT1_YqgM_like |
| conserved protein [Sulfolobus acidocaldarius DSM 639] | 71.2 | 71.2 | 98% | 5e-15 | 16.76% | AAY80607.1 | *saci_1262* | *essential* | MurG |
| hypothetical protein Saci_1921 [Sulfolobus acidocaldarius DSM 639] | 71.2 | 71.2 | 72% | 6e-15 | 13.33% | AAY81224.1 | *saci_1921* | MW069 BM-A055-B003 | RfaB |
| RecName: Full=Archaeal glycosylation protein 16 [Sulfolobus acidocaldarius DSM 639] | 65.9 | 65.9 | 75% | 3e-13 | 9.90% | Q4JAK2.1 | *saci_0807* | MW043 BM-A110-B008 | Archaeal glycosylation protein 16 |
| partially conserved Archaeal protein [Sulfolobus acidocaldarius DSM 639] | 61.2 | 61.2 | 96% | 1e-11 | 12.76% | AAY81207.1 | *saci_1904* | MW077 BM-A050-B002 | Cell wall/membrane/envelope biogenesis |
| glycosyl transferase [Sulfolobus acidocaldarius DSM 639] | 59.7 | 98.2 | 74% | 5e-11 | 14.02% | AAY80548.1 | *saci_1201* | MW053 BM-A150-B1 | Glycogen synthase |
| glycosyl transferase group 1 [Sulfolobus acidocaldarius DSM 639] | 47.8 | 47.8 | 77% | 3e-07 | 12.38% | AAY81133.1 | *saci_1827* | MW042 BM-A280-B001 | GT1_Trehalose_phosphorylase |
| gylcosyl transferase group 1 protein [Sulfolobus acidocaldarius DSM 639] | 45.8 | 45.8 | 49% | 1e-06 | 16.59% | AAY80595.1 | *saci_1249* | MW054 BM-A155-B001 | GT1_YqgM_like |
| conserved protein [Sulfolobus acidocaldarius DSM 639] | 42.4 | 42.4 | 34% | 1e-05 | 18.11% | AAY80235.1 | *saci_0869* | *essential* | Glycosyltransferase involved in cell wall biosynthesis |
| conserved protein [Sulfolobus acidocaldarius DSM 639] | 42.4 | 42.4 | 58% | 2e-05 | 9.91% | AAY81219.1 | *saci_1916* | MW066 BM-A024-B001 | |
| RecName: Full=UDP-sulfoquinovose synthase [Sulfolobus acidocaldarius DSM 639] | 40.4 | 40.4 | 26% | 7e-05 | 19.39% | Q4JBJ3.1 | *saci_0423* | MW039 BM-A083-B008 | Agl3 UDP-sulfoquinovose synthase |
| hypothetical protein Saci_1923 [Sulfolobus acidocaldarius DSM 639] | 39.3 | 39.3 | 34% | 1e-04 | 11.72% | AAY81226.1 | *saci_1923* | n.d. | |
| conserved protein [Sulfolobus acidocaldarius DSM 639] | 38.5 | 38.5 | 78% | 3e-04 | 11.85% | AAY81225.1 | *saci_1922* | MW070 BM-A060-B002 | |
| **ALG13** NP_011468.1 N-acetylglucosaminyldiphosphodolichol N-acetylglucosaminyltransferase [*Saccharomyces cerevisiae* S288C] | | | | | | | | | |
| conserved protein [Sulfolobus acidocaldarius DSM 639] | 43.9 | 43.9 | 58% | 1e-06 | 17.36% | AAY80607.1 | *saci_1262* | | MurG |
| **ALG14** NP_009626.1 N-acetylglucosaminyldiphosphodolichol N-acetylglucosaminyltransferase [*Saccharomyces cerevisiae* S288C] | | | | | | | | | |
| | No homolog above the standard Delta-Blast e-value threshold of 0.05 | | | | | | | | |
| **Alg14-13 fusion** (NP_009626.1+ NP_011468.1) | | | | | | | | | |
| conserved protein [Sulfolobus acidocaldarius DSM 639] | 59.7 | 59.7 | 73% | 4e-11 | 16.92% | AAY80607.1 | *saci_1262* | | |

**Table S2: Structural alignment modeling of Agl24 revealed a conservation of fold to the available structures of MurG and Alg13.** Structural modelling was performed by SWISS-MODEL (Waterhouse et al., 2018), using either the full length Agl24 sequence or only the c-terminal Alg13-like part. Identified PDB numbers as well as the SWISS_MODLE results, *e.g.* Global Model Quality Estimation (GMQE), quaternary structure quality estimate, and sequence identity are shown.

| PDB | GMQE: | QSQE: | Identity | X-ray | Reference |
|---|---|---|---|---|---|
| Full Agl24 sequence | | | | | |
| 3s2U MurG | 0.51 | - | 17.2 | 2.2Å | (Brown et al., 2013) |
| 1f0k MurG | 0.55 | 0,31 | 17.36 | 1.9Å | (Ha et al., 2000); |
| 1nlm MurG | 0.55 | 0,20 | 17.36 | 2.5Å | (Hu et al., 2003) |
| c-terminal Agl13-like part of Agl24 | | | | | |
| 2ks6 Alg13 | 0.56 | | 20.53 | NMR | (Raman et al., 2010) |
| 2jzc Alg13 | 0.56 | - | 19.87 | NMR | (Wang et al., 2008) |
| 1f0k MurG | 0.46 | 0.17 | 19.70 | 1.9Å | (Ha et al., 2000); |
| 1nlm MurG | 0.46 | - | 19.70 | 2.5Å | (Hu et al., 2003) |

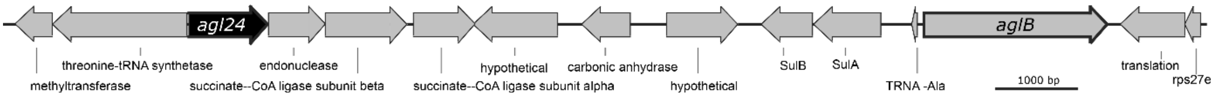


**Figure S1: Physical map of the gene region adjacent to *agl24 and aglB* of *S. acidocaldarius*.** Illustrated are the genes *Saci1260* until *Saci1276*. The gene *agl24* (*saci1262,* SACI_RS06030), displayed in black, which is annotated to encode a polysaccharide biosynthesis protein. The gene *aglB* (*saci1274,* SACI_RS06085), displayed with bold border, encodes the N-oligosaccharyltransferase, catalysing the transfer of the lipid-linked N-glycan onto the target protein.

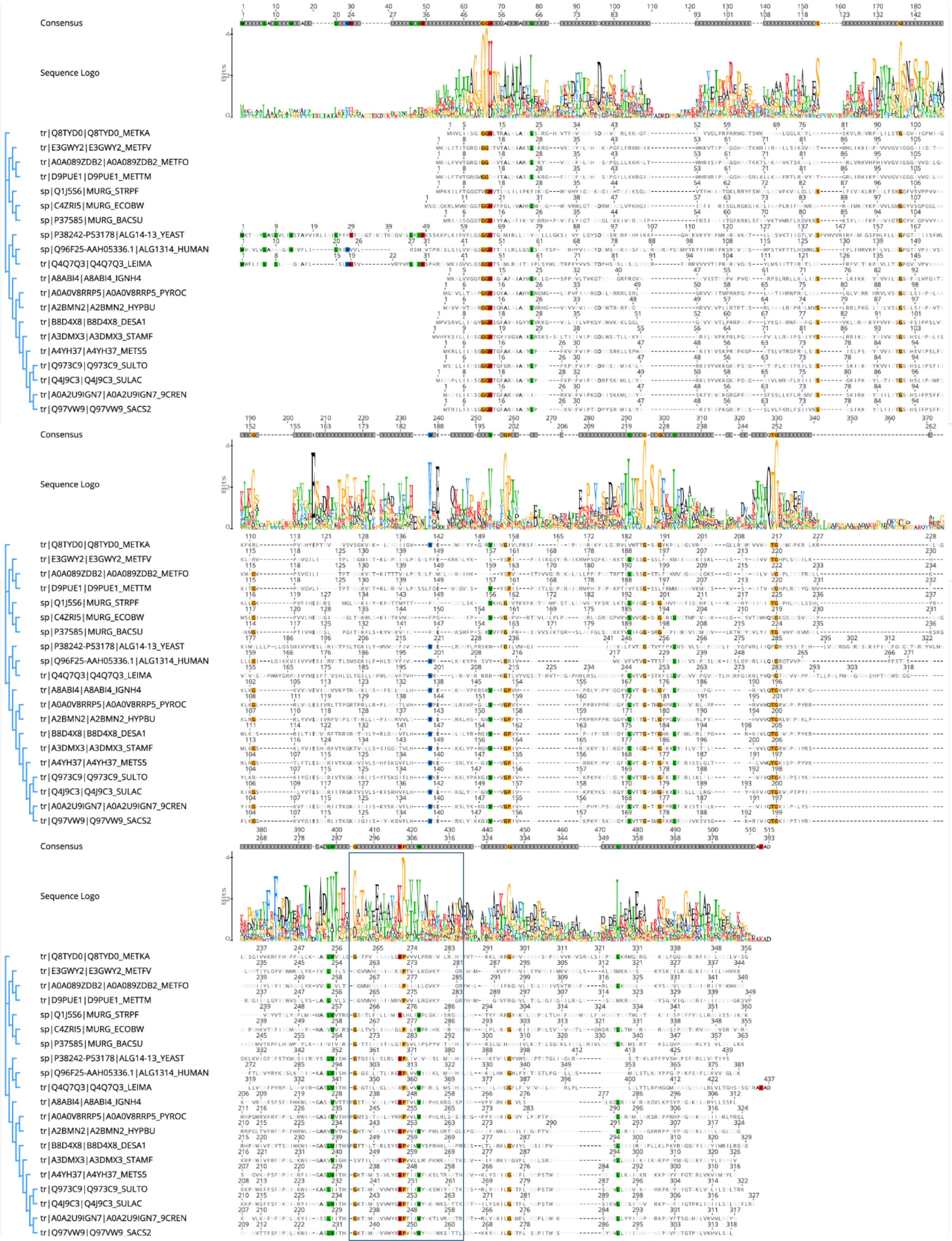

**Figure S2:** Protein sequences were aligned with the Clustal Omega (Sievers and Higgins, 2018). Sequences derived from i) three bacterial MurG: *E.coli* (C4ZRI5), *Streptococcus pyogenes* (Q1J5S6), and *Bacillus subtilis* (P37585); ii) eukaryal Alg14-13: *Leishmania major* (Q4Q7Q3), an artificial Alg14-13 fusion from *Saccharomyces cerevisiae* (P38242-P53178) and *Homo sapiens* (Q96F25-Q9NP73-2); iii) the crenarchaeal Agl24 homologs: *Ignicoccus hospitalis* (A8ABI4), *Pyrodictium occultum* (A0A0V8RRP5), *Hyperthermus butylicus* (A2BMN2), *Desulfurococcus amylolyticus* (B8D4X8), *Staphylothermus marinus* (A3DMX3), *Metallosphaera sedula* (A4YH37), *Sulfurisphaera tokodaii* (Q973C9), *Sulfolobus acidocaldarius* (Q4J9C3), *Acidianus brierleyi* (A0A2U9IGN7), and *Saccharolobus solfataricus* (Q97VW9). Selected sequences from pseudomurein producing Euryarchaea with higher sequence similarity to the MurG are aligned: *Methanopyrus kandleri* (Q8TYD0), *Methanothermus fervidus* (E3GWY2), *Methanobacterium formicicum* (A0A089ZDB2), *Methanothermobacter marburgensis* (D9PUE1). Conserved amino acids (65% threshold) are highlighted in color corresponding to their chemical properties of the amino acids: green polar amino acids (G, S, T, Y, C, Q, N), blue basic (K, R, H), red acidic (D, E), and black hydrophobic amino acids (A, V, L, I, P, W, F, M). End of the Agl14-like domain and start of Alg13-like domains are indicated by the change from dark to light background color. Conserved amino acids (65% threshold) are highlighted in color. Weblogo from the sequence aliment shown above. Height of each bar correspond to the observed frequency. **Boxed**: area of the sugar donor binding site, in *E. coli* MurG the amino acids $A_{264}$, $L_{265}$, $T_{266}$, $E_{269}$, $Q_{288}$, and $Q_{289}$ have been to interact with UDP-GlcNAc.

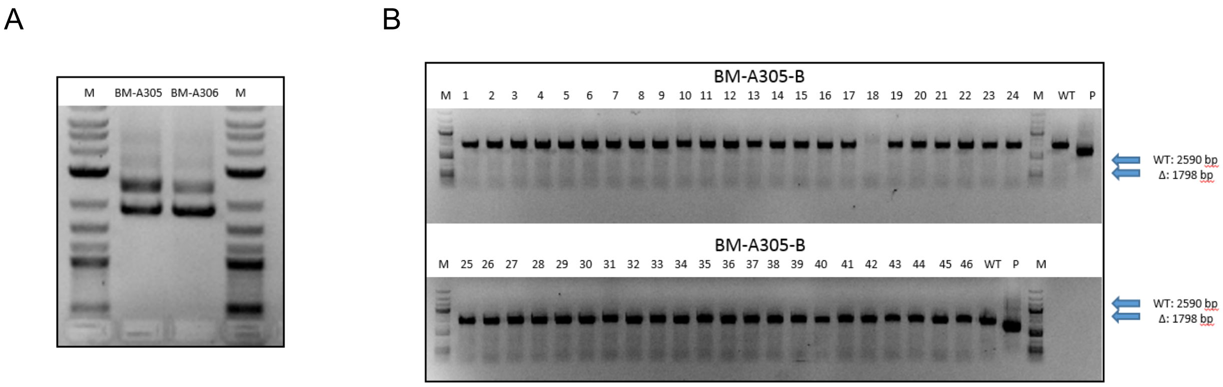


**Figure S3: Confirmation of the integration and segregation of the *ag23* deletion plasmid pSVA1312 in *S. acidocaldarius* MW001 A)** The integration of the *Agl24* deletion plasmid pSVA1312 in *S. acidocaldarius* MW001 (first selection) was monitored by PCR using the out primers of the upstream and downstream region of *Agl24* and the genomic DNA from two first selection colonies incorporate the plasmid (BM-A305 and BM-A-306). DNA from the background strain MW001 and the plasmid pSVA1312 were used as control, showing a PCR fragment corresponding to the flanking region including or excluding the *Agl24* gene, respectively. **B)** The segregation of pSVA1312 (second selection) was confirmed by PCR using the outer primers of the flanking region of *Agl24* and the genomic DNA from second selection colonies. All PCR fragments gained from genomic DNA of second selection colonies correspond to the full length *Agl24* gene (2590 bp), while a deletion would result in a 1798 bp PCR fragment.

A

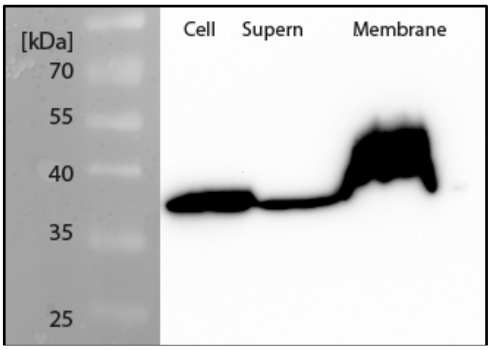


B

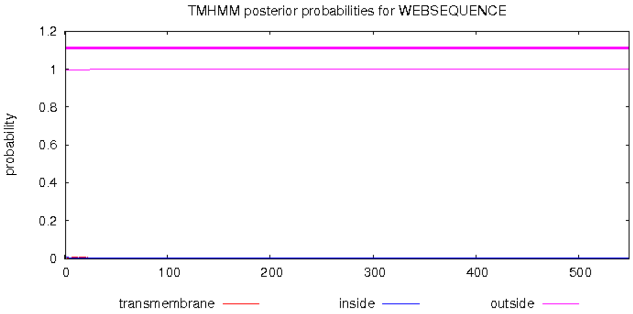


C

```
MIDNPLLIIASGGGHTGFARAIAEYLPFKPDFVIPENDRFSKDMLLDYARKLYYVKKGKDPGQGNIVLMRNFLKIIIESGKIPKYLATIATGSN
----EEEEEEE-----HHHHHHHHHHH----EEEE----HHHHHH------EEEE--------HHHHHHHHHHHHHHHHHHHHH---EEEE----
HSLIPAMFQKIKGSALYVTESQDRIITRGKTVSVLSKFSRHVFLHWNEQKGLYDNGVVVGPIVEKPKYKSENKGYILVTTGSMGFKKLFDSLLN
--HHHHHHHHH----EEEEE---EEEE---HHHHH-----EEE------HHH----------------------EEEEEE----HHHHHHHHHH
LRGNYKFVIQTGKVDPTPYIEKKPDWSFFSFDKDIERYIANAELVITHQGKTAMESVVMYGKPTIIVYNKDWKSATTKQDTILYSKILGATFLD
-----EEEEEE-----HHHHH----EEEEE----HHHHHHHH-HEEE---HHHHHHHHHH---EEEEE--------HHHHHHHHHHHH--EEEE
DPSTWDSIKVLEDSIQNVRKPNQFEIGTPKLIDIILSELAEFLRE
------HHHHHHHHHHHHH---HHHHHHHHHHHHHHHHHHHHH--
```

**Figure S4: Localization of Agl24 in *S. acidocaldarius*. A**. Anti-His immunoblot of samples from *S. acidocaldarius* BM-A724 expressing Agl24 with C-terminal His- and Strep-Tag reveal that the majority of Agl24 is found in the membrane fraction. Cell: total cell pellet, **B.** Prediction of transmembrane helices in Agl24, by the TMHMM Server v. 2.0 (http://www.cbs.dtu.dk/services/TMHMM/) confirmed the lack of any transmembrane domain. **C** Secondary protein structure prediction by the Jpred Server (Drozdetskiy et al., 2015).

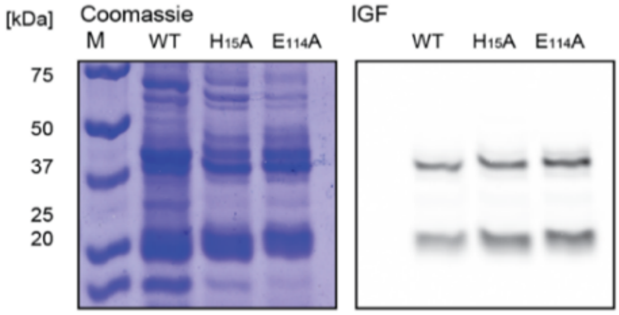


**Figure S5**: SDS-PAGE of the purified Agl24-WT-GFP, mutants Agl24-$H_{15}$A-GFP, and Agl24-$E_{114}$A-GFP from *E.coli*, stained with Coomassie Brilliant Blue or visualized by in gel fluorescents (IGF).

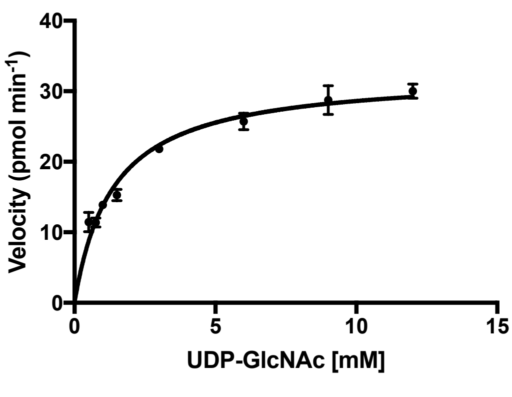


| | |
|---|---|
| Michaelis-Menten | |
| Best-fit values | |
| Vmax | 32.53 |
| Km | 1.366 |
| Std. Error | |
| Vmax | 0.8961 |
| Km | 0.1282 |
| 95% CI (profile likelihood) | |
| Vmax | 30.68 to 34.53 |
| Km | 1.112 to 1.675 |
| Goodness of Fit | |
| Degrees of Freedom | 21 |
| R square | 0.9548 |
| Absolute Sum of Squares | 56.05 |
| Sy.x | 1.634 |
| Constraints | |
| Km | Km > 0 |
| Number of points | |
| # of X values | 24 |
| # Y values analyzed | 23 |

**Figure S6: Agl24 $K_m^{app}$ for UDP-GlcNAc determination**

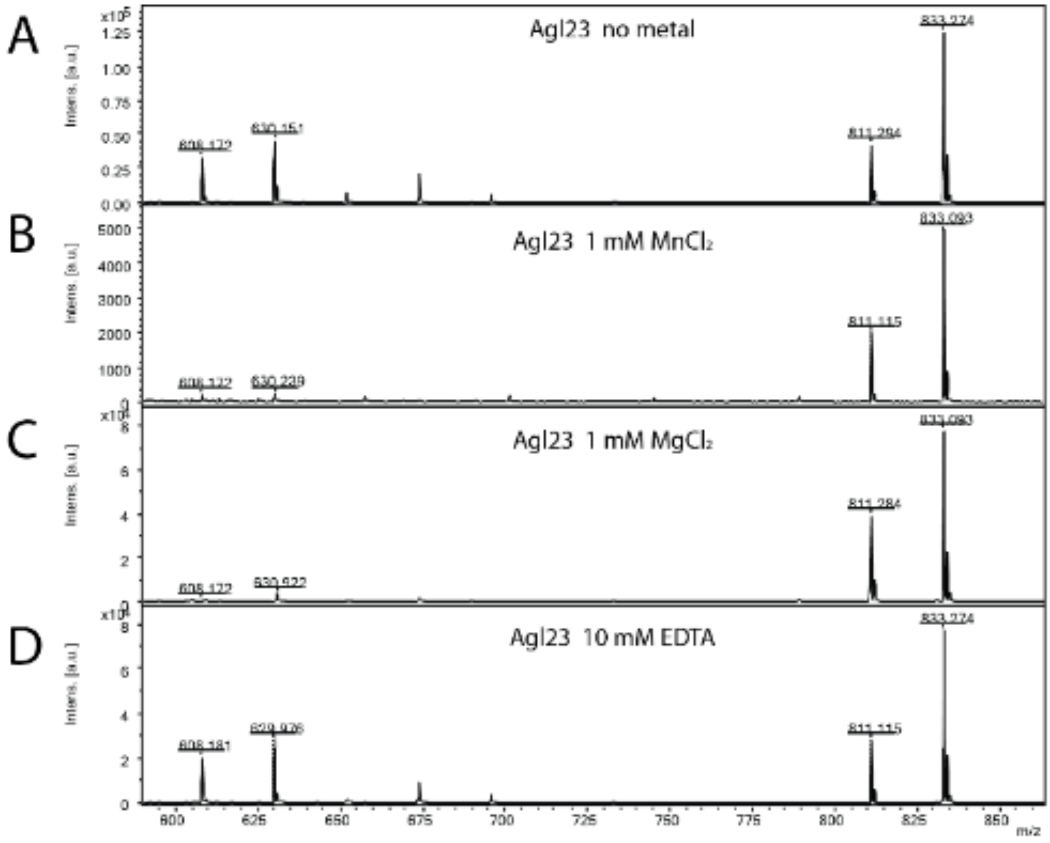


**Figure S7: MALDI-MS spectra of the *in vitro* Agl24 reaction to assess the metal dependency for the specificity activity.** Spectra obtained from the purified enzymatic reaction mix with acceptor-1, UDP-GlcNAc and Agl24-GFP without addition of metal cations (**A**), with the addition of 1 mM $MnCl_2$ (**B**), 1 mM $MgCl_2$ (**C**), or 10 mM EDTA (**D**). In all cases the acceptor-1 (608 m/z [M-1H+2Na] and 630 m/z [M-2H+3Na]) was converted to the product (811 m/z [M-1H+2Na] and 833 m/z [M-2H+3Na]), indicating the metal cations are not required for enzymatic activity.

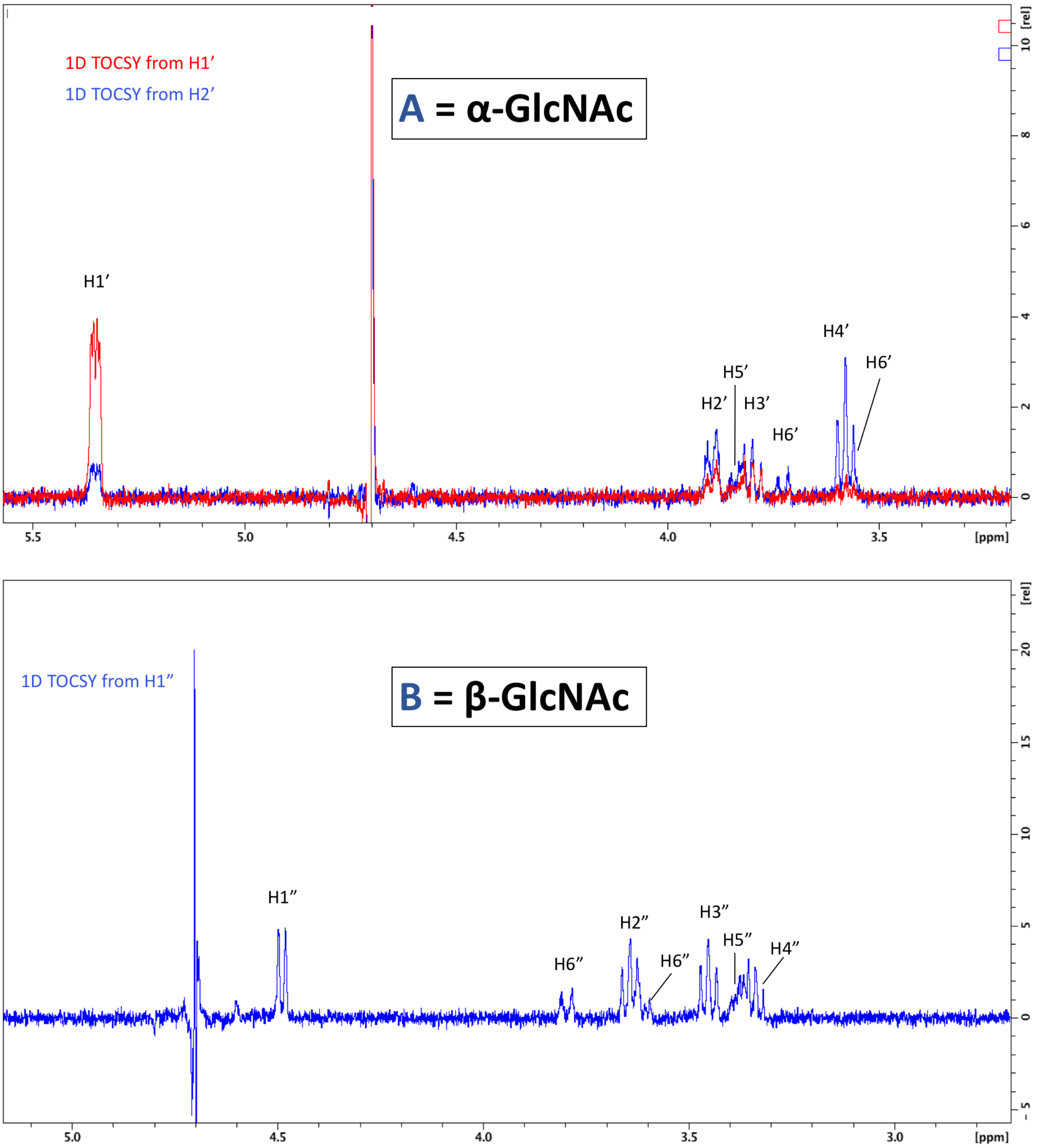


**Figure S8: 1D TOCSY correlation spectra**. To visualize all protons of the $\alpha$-linked GlcNAc residue, irradiation was set for the shift of H1’ and H2’ (Top). To visualize protons of the $\beta$-linked GlcNAc, irradiation of H1” was sufficient to see all protons (Bottom).

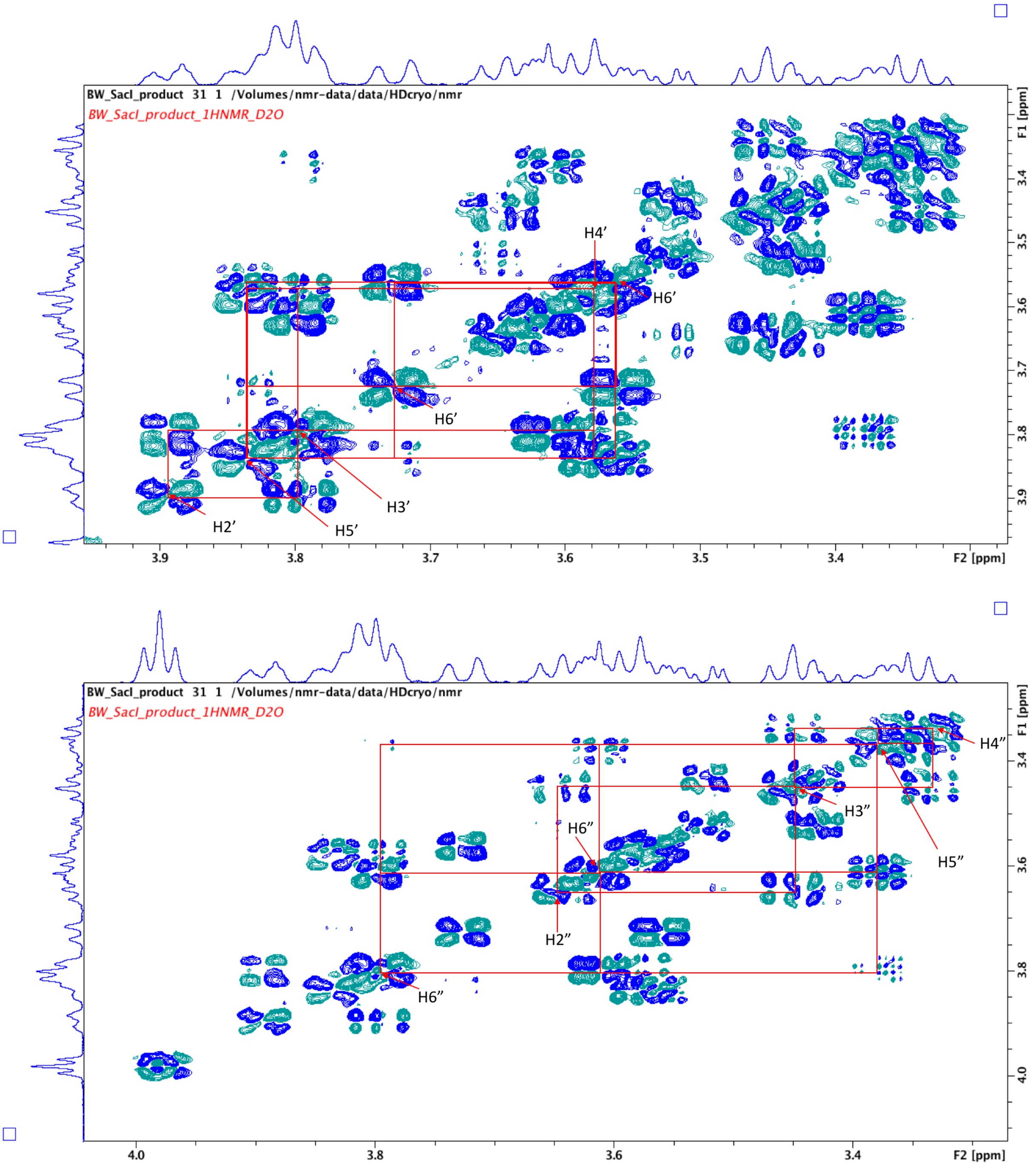


**Figure S9: 2D COSY spectra used to assign proton signals arising from α-GlcNAc (Top) and β-GlcNAc (Bottom).**

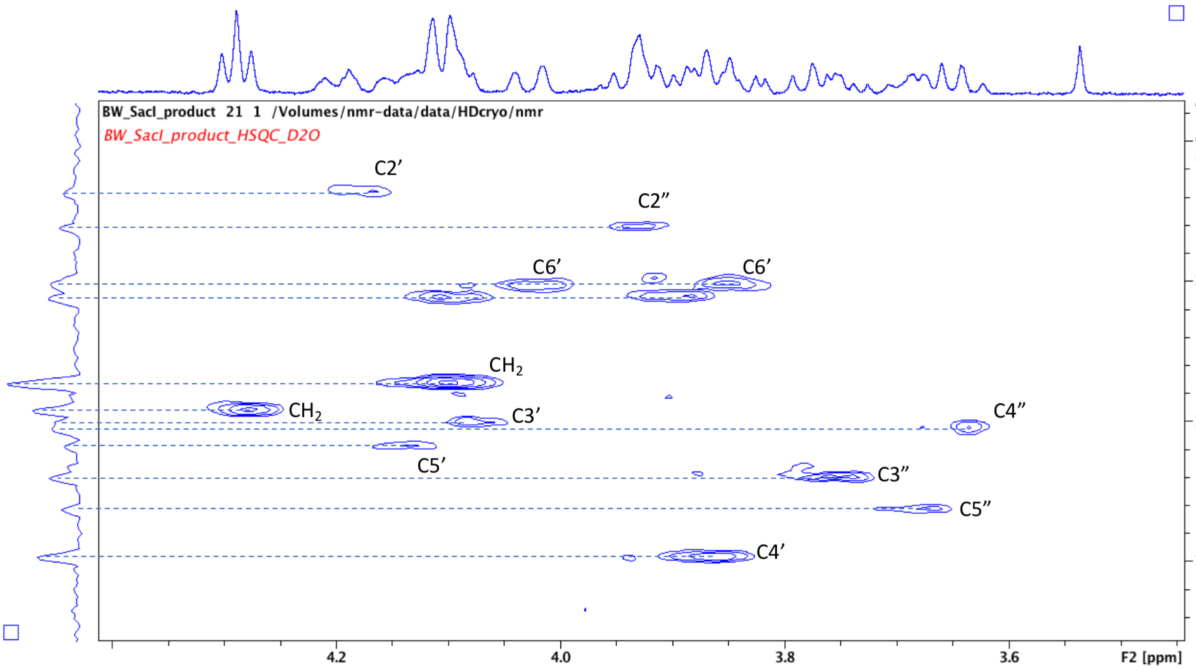


**Figure S10: 2D HSQC spectra used to assign carbon signals arising from both α- and β-linked GlcNAc residues.** Note the signal for C4’ is significantly shifted (79.6 ppm) due to the presence of the glycosidic linkage at this position.

**Figure S11: Universal distribution of Agl24 homologs in Crenarchaeota, white the exception of the Order Thermoproteaces**. The BLAST analyses (https://blast.ncbi.nlm.nih.gov) of Agl24 (Q4J9C3) with the restriction to Crenarchaeota (Taxid:28889; 126 genomes), revealed 100 sequences (30-100% sequence identity), lacking any homology within the 27 genomes of the Order Thermoproteaces. The orders of Crenarchaeaota are background colored: Fervidicoccales (green), Acidilobales (blue), Desulfurococcales (orange), Sulfolobales (yellow), Thermoproteales (red).

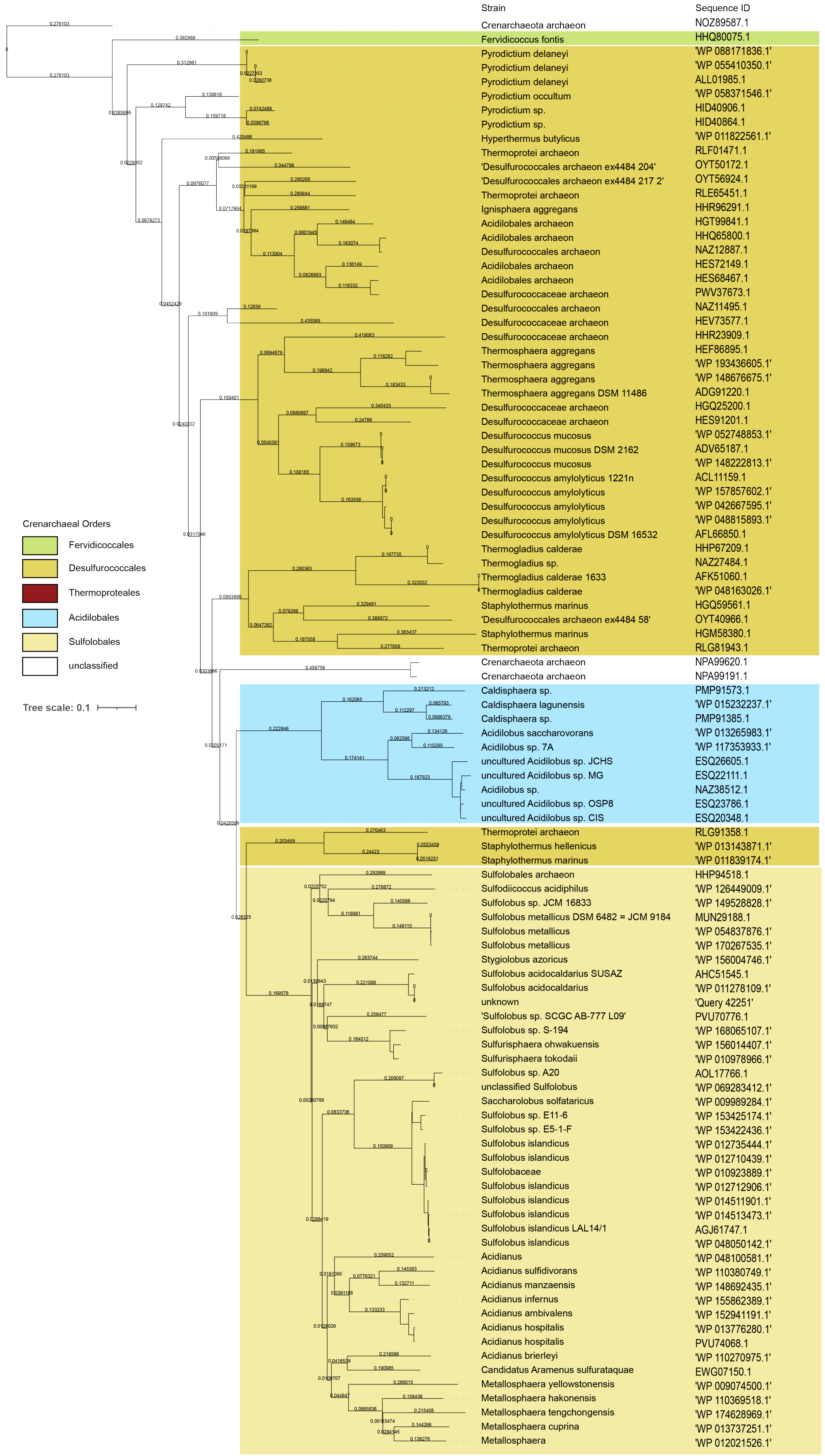

**Table S3: Strains and plasmids used in this study.**

| Strains | Genotype | Reference |
|---|---|---|
| ***E. coli*** | | |
| DH5α | F– ϕ80lacZΔM15 Δ(lacZYA-argF)U169 recA1 endA1 hsdR17(rK–, mK+) phoA supE44 λ– thi-1 gyrA96 relA1 | (Taylor et al., 1993) |
| BL21 (DE3) | $F^-$ *ompT* *hsd*$S_B$ ($r_B^-$, $m_B^-$) *gal dcm* (DE3) | (Studier and Moffatt, 1986) |
| ER1821 | F- glnV44 e14-(McrA-) rfbD1? relA1? endA1 spoT1? thi-1 Δ(mcrC-mrr)114::IS10, MM294 background | (Meselson and Yuan, 1968) |
| *Sulfolobus acidocladarius* | | |
| MW001 | *S. acidocaldarius* DSM 639, Δ*pyr*E | (Wagner et al., 2012) |
| BM-A305 | *S. acidocaldarius MW001* with integrated deletion plasmid pSVA1312 incl. *pyr*EF | *This study* |
| BM-A306 | *S. acidocaldarius MW001* with integrated deletion plasmid pSVA1312 incl. *pyr*EF | *This study* |
| BM-A719 | *S. acidocaldarius MW001* with pSVA1337 ($MAL_{promotor}$ *saci1262,* C-terminal Strep- and His-Tag) | *This study* |
| BM-A720 | *S. acidocaldarius MW001* with pSVA1339 ($MAL_{promotor}$ *saci1262*, N-terminal Strep- and His-Tag) | *This study* |
| BM-A724 | *S. acidocaldarius MW001* with pBM-0078 ($ARA_{promotor}$ *saci1262* C-terminal Strep- and His-Tag) | *This study* |
| **Plasmids for generating deletion mutants** | | |
| pSAV0407 | Gene targeting plasmid, pGEM-T Easy backbone, *pyrEF* cassette of *S. solfataricus* | (Wagner et al., 2012) |
| pSVA1312 | In-frame deletion of *Agl24* (*saci1262*) cloned into pSVA407 with *Apa*I and *Bam*HI | *This study* |
| pSVA3338 | Linear *saci1262*($_{short}$)-*pyrEF-saci1262* cut with *BamH*I and *Apa*I into pSVA407 | *This study* |
| **Expression plasmids for *S. acidocaldarius*** | | |
| pSVA1481 | *E. coli* vector with Ara-promoter and C-terminal Strep- and His-tag based on pGEM-T Easy backbone and pMZ1 cassette, pMZ: $Amp^r$, cloning vector containing replicon ColE1 (pBR322) and a 10X His-Strep tag sequence at the 3' end of the MC | Wagner & Albers |
| pSVA1518 | Expression vector with $ARA_{promotor}$ and c-terminal Strep-$His_{10}$ tag | Wagner & Albers |
| pSVA1431 | Expression vector with $MAL_{promotor}$ and c-terminal Strep-$His_{10}$ tag | Wagner & Albers |
| pSVA2301 | Expression vector with $MAL_{promotor}$ and N-terminal $His_{10}$- Strep- tag | Wagner & Albers |
| pSVA1336 | *Agl24* cloned into pSVA1481 ($ARA_{promotor}$)with a C-terminal Strep- and His-tag with *Nco*I and *Pst*I | *This study* |
| pSVA1337 | *Agl24* with c-terminal Strep- and His-Tag derived from pSAV1336 subcloned into pSVA1431 ($MAL_{promotor}$) with *NcoI* and *EagI* | *This study* |
| pSVA1339 | Agl24 cloned into pSVA2301 with N-terminal $His_{10}$ Strep - TEV into pSVA2301 ($MAL_{promotor}$) with *NcoI* and *NotI* (Primer 4180 and 4181*)* | *This study* |
| pBM-0078 | Ara promotor saci1262 with C-terminal Strep- and His-tag, cloned with *NcoI ApaI* | *This study* |
| **Expression plasmids for *E. coli*** | | |
| *p*Waldo | T7 expression vector containing TEV site-GFP-$His_8$ | *(Waldo et al., 1999)* |
| pHD0499 | pWaldo containg *Agl24*-GFP-$His_8$ cloned with *Xho*I and *Kpn*I | *This study* |
| pHD0554 | pWaldo containg *Agl24*-$H_{14}$A-GFP-$His_8$ cloned with *Xho*I and *Kpn*I | *This study* |
| pHD0586 | pWaldo containg *Agl24*-$E_{114}$A-GFP-$His_8$ cloned with *Xho*I and *Kpn*I | *This study* |

**Table S4: Primers used in this study.**

| Primer | Sequence (5´-3´) | Restriction Site |
|---|---|---|
| ***ΔAgl24*** | | |
| 4168 | ACTA GGGCCC GGTCTTGTGCTTAAATTCACTCTGACATC | *Apa*I |
| 4163 | GTC TAA AAA TGT GGC TCC TCC ACT GGC GAT AAT CAG TAA TGG GTT GTC | |
| 4164 | ATC GCC AGT GGA GGA GCC ACA TTT TTA GAC GAT CCC TCG ACC TGG GAC 4165 | |
| 4165 | CGCCGA GGATCC CAC AAA TAT ATT CTC CCA AGA GTC TGG C | *BamH*I |
| **pSVA3338 Linear fragment *Agl24*$_{up}$*-pyrEF-Agl24*$_{down}$** | | |
| 4168 | ACTA GGGCCC GGTCTTGTGCTTAAATTCACTCTGACATC | *ApaI* |
| 6336 | GTA ACA TAT CTT TGC TAA ATC GGT CAT TTT CGG GTA TTA C | |
| 4115 | GTAATACCCGAAAATGACCGATTTAGCAAAGATATGTTACTCGATTACGCTAGAAAA TTTGAGCAGTTCTAGTACTTGGCTCAAAGAATG | |
| 4116 | GGT GTA CCT ATT TCG AAT GGT TTA GGT TTT CTA ACA TTT TGG ATA CTA TCT CAC TAA TTT CAT TTT TTC CTA AAA ATT GCT CCT TTA CAT TTC | |
| 6337 | GTATCCAAAATGTTAGAAAACCTAACCAATTCGAAATAGGTACACC | |
| 6338 | CGCCGA GGATCC GAA TGC CCA ATT TTT TCA TTT CTG CAA TAG TCT GTT C | *BamHI* |
| ***S. acidocaldarius* Expression** | | |
| pSVA1336 | | |
| 4176 | CTGCA CCATGGG T ATCGACAACCCATTACTGATTATCGCCAGTGG | *NcoI* |
| 4177 | CCGTT CTGCAG C TCT AAG AAA TTC TGC TAA TTC ACT TAA AAT TAT ATC | *PstII* |
| pSVA1339 | | |
| 4180 | CTGCA CCATGGG TCGACAACCCATTACTGATTATCGCCAGTGG | *NcoI* |
| 4181 | CCGTT GCGGCCGC TTA TCT AAG AAA TTC TGC TAA TTC ACT TAA AAT TAT ATC | *NotI* |
| ***E. coli* Expression of *agl24*** | | |
| A596 | *saci1262*-fwd-*XhoI* AGGAGA CTCGAG ATGATCGACAACCCATTACTGATTATCGCC | *XhoI* |
| A597 | *saci1262*-rev-*KpnI* GATCCA GGTACC TCTCTAAGAAATTCTGCTAATTCACTTAAAATTATATC | *KpnI* |
| A684 | *saci1262*_$H_{14}$A-*XhoI*-fwd AGGAGACTCGAGATGATCGACAACCCATTACTGATTATCGCCAGTGGAGGAGGGGCTACTGGCTTCGCTAGAGCTATTGC | *XhoI* |
| A685 | *saci1262*_$E_{114}$A-fwd GTGCACTTTATGTAACAGCTAGCCAAGACAGAATTATTAC | |
| A686 | *saci1262*_$E_{114}$A-rev GTAATAATTCTGTCTTGGCTAGCTGTTACATAAAGTGCAC | |

**Table S4: Overview of the properties of the archaeal, eukaryal and bacterial N-glycosylation.** Highlighted in grey background color are the mammalian orthologs of the non-catalytic OST subunits found in *Saccharomyces cerevisiae.* *based on sequence analyses of metagenome-assembled genomes of Asgardarchaeota (Zaremba-Niedzwiedzka et al., 2017), **not present in STT3 proteins from organisms that express single subunit OSTs. ***chitobiose is not present in Thermoproteales, but showing an modified version: (GlcA(NAc)2-β-1,4-(Glc(NAc)2

| Domain / Component | **Eukarya** | **Archaea** | | | **Bacteria** |
|---|---|---|---|---|---|
| | | Asgardarchaeota | Crenarchaeota | Euryarchaeota | |
| Lipid carrier | Dol | Dol | Dol | Dol | Und |
| Linkage between N-glycan and lipid carrier | PP | ? | PP | P | PP |
| Nucleotide activated sugar donor | yes | ? | yes | yes | yes |
| Lipid activated sugar donor | yes | ? | yes | yes | no |
| Oligosaccharyltransferase | Stt3/ STT3A/STT3B | AglB (Stt3-like) | AglB (Stt3-like) | AglB (PglB-like) | PglB |
| DK motif (DXXKXXX(M/I) | yes | yes | yes | yes | |
| DKi motif (D/E< >KXXXM/I/P) | | | | yes | |
| MI motif (MXXIXXX(I/V/W) | | | | yes | yes |
| double sequon DNXTZNXS/T | yes** | yes | yes | | |
| Non-catalytic OST subunits | Ost1/RPN1<br>Ost5/TMEM258<br>Swp1/RPN2<br>Wbp1/DDOST<br>Ost2/DAD1<br>Ost4 / Ost4<br>Ost6/TUSC3<br>or Ost3/MAGT1 | RPN1(ribophorin-1)*<br>Ost5*<br><br>Wbp1*<br><br><br>Ost6-like/Ost3-like* | n.d. | n.d. | n.d. |
| First enzymes in the glycosylation | Alg7<br>Agl14/13 | ? | AglH<br>Agl24 | diverse | PglC<br>PglA |
| N-glycan linking sugar(s) | Chitobiose<br>(GlcNAc- β 1,4- GlcNAc -β1-) | ? | Chitobiose*** | diverse | GalNAc-α1,3-*Bac*-β1 |
| N-glycan structure | tri-branched | ? | di- and tri-branched | linear, di-, tri-branched | linear |

**Supplementary Data 1:**

Seed sequences, homology searches, preliminary phylogenies, final datasets (individual homologs), final datasets (full and trimmed alignments), and final phylogenies for the phylogenetic analysis are collected in a zip file.
